# Estimating avian wingspan from wing length: an order-level regression approach

**DOI:** 10.64898/2026.04.27.720532

**Authors:** M.R. Fox, E.A. Brunton, J.M. Shephard

## Abstract

Wingspan is a key parameter in avian collision risk models but is rarely measured systematically, meaning for many species, collision risk remains largely unquantified due to a lack of morphological and flight data required to parameterise models. Folded wing length scales predictably with wingspan and is available for all extant bird species, yet no widely available method has been published for predicting wingspan from wing length; here we compiled a global wingspan dataset for 1,442 species across 25 orders and fitted order-level linear regression models predicting wingspan from wing length. Order-level models performed well, with 85% achieving R^2^ ≥ 0.95 (global model R^2^ 0.96); cross-validation identified that model predictions generally fell within the intraspecific variation in wingspan, resulting in wingspan estimates for 9,194 species. Our models and accompanying wingspan data provide a transparent, reproducible, and validated method for generating species-level wingspan estimates from widely available wing length data, with direct applications to collision risk modelling, flight performance research, and macroecological analyses of avian morphology.

## INTRODUCTION

Wing morphology is a primary determinant of avian flight performance shaping foraging behaviour, flight speed, dispersal ability, and energetic costs across species (Alerstam et al., 2007; Fu et al., 2023; Pennycuick, 2008). Wing length is among the most widely measured avian morphological traits and is compiled globally for all extant bird species in AVONET (Tobias et al., 2022). Wingspan, the tip-to-tip distance of fully extended wings, is considerably less accessible. Unlike wing length, which is routinely measured from a bird in hand, wingspan data are more difficult and time-consuming to obtain requiring wings to be fully spread or multiple measurements. Consequently, wingspan is rarely recorded alongside other morphological traits (Fu et al., 2023; Shiomi et al., 2025). *BirdWingData* (Shiomi et al., 2025) is the most comprehensive wingspan database currently available and covers 856 species, only c. 8% of extant species, leaving a considerable knowledge gap, which has significant practical consequences. For example, wingspan determines wing loading and aspect ratio, underpinning estimates of flight speed and energetic costs that are relevant to migration and behavioural research (Grilli et al., 2017; KleinHeerenbrink et al., 2015; Pennycuick, 2008).

Critically, wingspan is a key input for many collision risk models (CRM) that assess bird collision risk with wind turbines, a research area that has gained increased attention in recent decades (Cook et al., 2025; Masden et al., 2021). Empirical wingspan data are absent for many species for which such assessments are required. Collision risk varies considerably among species, with vulnerability driven by species-specific physical and behavioural traits including wing morphology, life history characteristics and flight height and type (Biasotto et al., 2022; Gémard et al., 2025; Thaxter et al., 2017). Variation in species traits means that some taxonomic groups face greater collision risk and population-level consequences than others. In particular, raptors (Accipitriformes) are consistently identified as the highest-priority group for collision risk across North America and Europe, as well as migratory species and seabirds, with population-level consequences most likely for long-lived, slow-reproducing species (Beston et al., 2016; Desholm, 2009; Estellés-Domingo & López-López, 2025; Thaxter et al., 2017). In Australia, cockatoos and migratory parrots are also ranked among the highest for potential impacts of onshore and offshore wind energy (Reid et al., 2025; Reid et al., 2023). Seabirds, and particularly Procellariiformes, are of growing concern as offshore wind energy expands into the Southern Hemisphere. For many of these species, collision risk remains largely unquantified due to a lack of morphological and flight data required to parameterise CRMs (Furness et al., 2013; Miller et al., 2025; Reid et al., 2025). The potential for birds to collide with wind turbines is typically assessed in environmental impact assessments (EIA) through the use of a CRM (Masden & Cook, 2016). Cook et al. (2025) reviewed 52 different types of CRMs, and while CRMs vary in their input data requirements, several inputs are consistent between models (Cook et al., 2025). Of the models reviewed by Cook et al. (2025), 58% require wingspan as input data, and 62% require flight speed, which can be calculated using morphological data including wingspan (Klein Heerenbrink, 2023; Pennycuick, 2008). Widely used CRMs and those frequently used in environmental impact assessments use wingspan as a key model input, yet it is poorly represented in available trait databases (Band & Band, 2012; Band et al., 2007; Smales et al., 2013).

Wing length and wingspan are both measures of wing size and are expected to scale allometrically (Nudds, 2007), suggesting that wingspan may be readily predicted from the more widely available folded wing length (hereafter wing length) measure (Biasotto et al., 2022; BirdLife International, 2025). Some variation in wingspan–wing length relationship would be expected across bird orders, reflecting evolutionary differences in wing morphology associated with flight behaviour (Sheard et al., 2020; Sullivan et al., 2019). For example, the disproportionately long wings of dynamic-soaring Procellariiformes are likely to be morphologically dissimilar to the compact wings of aerial insectivores. However, no widely available method has been published for predicting wingspan from wing length, meaning that researchers and practitioners requiring wingspan data for species where it has not been directly measured, currently have no validated approach for doing so.

Here we compile wingspan data from *BirdWingData* (Shiomi et al., 2025), Birds of the World (Cornell Lab of Ornithology, 2026), and the Handbook of Australian New Zealand and Antarctic Birds (HANZAB) (BirdLife Australia, 2023), and combine these with wing length data from AVONET. Using these data we developed and validated order-level linear regression models for predicting avian wingspan from wing length and applied these models to predict wingspan data for 9,194 flighted bird species. We discuss the utility of having more widely available wingspan data and the broader value of having wingspan estimates available at a global scale.

## METHODS

We extracted data from *BirdWingData* (Shiomi et al., 2025), which provides measured wingspan data for a subset of species. *BirdWingData* has two broad categories of wingspan data based on the measurement method, we omitted all B measures which exclude the width of the body between the two wings, thereby underestimating wingspan (Shiomi et al., 2025). We then searched the HANZAB via BirdLife Australia (BirdLife Australia, 2023), and species-specific accounts in Birds of the World (Cornell Lab of Ornithology, 2026), using the keyword “wingspan”.

Searches were prioritised toward orders previously identified as having elevated collision risk (i.e., raptors, seabirds, waterbirds), including Accipitriformes, Falconiformes, Procellariiformes, Suliformes, Pelecaniformes, Ciconiiformes, and Charadriiformes. To achieve a representative sample, we searched every flighted order, aiming for each family to be represented in the search effort, except for Passeriformes due to the large number of families. For Birds of the World, keyword searches were conducted on the main landing page of each species account and, where available, under the “Plumage, Molts, and Structure” tab, which contained the subheading “Measurements”. Where a range was reported (e.g., 100–120 cm), both the minimum and maximum values were retained. We restricted our search to species-level entries, excluding subspecies accounts. Wing length data for all species were sourced from AVONET (Tobias et al., 2022), which provides standardised morphological measurements from museum specimens. All data were matched across sources using a hierarchical name-matching procedure to account for taxonomic discrepancies between the BirdLife, eBird, and BirdTree naming conventions used by each source.

To assess the relationship between wing length and wingspan, we fit linear regressions to the data. We used the reported mean for wing length data from AVONET and calculated mid-points for wingspan data across sources. We removed species belonging to flightless or poorly flighted orders (Struthioniformes, Sphenisciformes, Mesitornithiformes, Cariamiformes, Galliformes). We first fitted a global linear model (wingspan ∼ wing length) across all species with empirical wingspan data. We then fitted order-level linear models separately for each order with five or more species with wingspan data. To improve coverage for orders with fewer than five species, several small orders were grouped with functionally or phylogenetically similar larger orders prior to modelling. We followed IOC (International Ornithological Congress) taxonomy (Gill et al., 2025), and grouped Cathartiformes (New World vultures) with Accipitriformes, and split Caprimulgiformes into Caprimulgiformes (families: Caprimulgidae, Podargidae) and Apodiformes (Trochilidae, Apodidae, Aegothelidae, Hemiprocnidae). Phaethontiformes (tropicbirds) were grouped with Charadriiformes (shorebirds) based on visual inspection of the wing length–wingspan relationship; Phoenicopteriformes (flamingos) were grouped with Pelecaniformes, and Pterocliformes (sandgrouse) with Columbiformes (pigeons and doves). Gaviiformes (loons) and Podicipediformes (grebes) were combined into a single group given their functional similarity as aquatic diving birds and limited data availability (n = 9). We then plotted data and linear regression curves, using minimum and maximum values for wingspan and wing length to demonstrate intraspecific variation.

To evaluate model predictive accuracy, we conducted a 1,000-iteration repeated random sub-sampling cross-validation in which 20% of species within each order were randomly withheld in each iteration, with the remaining 80% used to fit order-level linear models. For each iteration, wingspan was predicted for the withheld species using the order-level model fitted on the training data. Prediction accuracy was quantified as the absolute percentage deviation between the predicted wingspan and the reported midpoint wingspan for each withheld species. To contextualise model prediction error against biological variation, we calculated the intraspecific range in wingspan for species with empirical minimum and maximum values reported in the source literature, expressed as the difference between the reported midpoint and the reported maximum (or minimum) wingspan as a percentage of the midpoint.

Using the order-level and global models, we predicted wingspans for all flighted bird species, excluding Galliformes, in AVONET lacking empirical wingspan data (n = 9,194). Predicted minimum and maximum wingspan values were derived by applying the order-specific median intraspecific range (%), for orders with fewer than five species with empirical range data the overall median (%) was used. All analyses were conducted in RStudio version 5.1 (R Core Team, 2025).

## RESULTS

We collated wingspan data for 1,442 species, representing approximately 13% of all extant bird species; the full dataset is available (Appendix S1). Wingspan data were sought for 3,568 species across HANZAB and Birds of the World combined; retrieval rates differed substantially between sources, with 79.1% of species searched yielding data in HANZAB compared to 11.2% in Birds of the World (Table 1). Several orders had considerable search effort but low retrieval (Table 1): Piciformes, Bucerotiformes, Columbiformes, and Strigiformes. In contrast, Falconiformes, Accipitriformes, Procellariiformes, and Suliformes were well represented (Table 1). Passeriformes, Psittaciformes, and Cuculiformes had lower representation overall; for these orders, the majority of records were sourced from HANZAB (Table 1).

**Table 1:** Summary of wingspan data collated and search effort by avian order. For each order, the number of species with wingspan data retrieved from BirdWingData (BWD; Shiomi et al., 2025; excluding species with only definition B data, which excludes the inter-wing body segment and therefore underestimates total wingspan), the Handbook of the Birds of Australia, Antarctica and New Zealand (HANZAB; BirdLife Australia, 2023), and Birds of the World (BOTW; Cornell Lab of Ornithology, 2026) are presented, along with the number of species searched in HANZAB and BOTW. The total number of unique species with wingspan data and the percentage of the order represented in the final dataset are also shown. Orders marked with † were largely excluded from search efforts and excluded from modelling due to flightlessness or atypical flight morphology. Searches were conducted using the keyword “wingspan” on species account landing pages and, where available, under the “Plumage, Molts, and Structure” tab in Birds of the World. BirdWingData was accessed as a database extract and species were not individually searched.

| Order | Species wingspan data retrieved |  |  |  | Number of species searched for wingspan data |  |  | Retrieval rate from BOTW & HANZAB | Total unique found | % of order available from search effort |
| --- | --- | --- | --- | --- | --- | --- | --- | --- | --- | --- |
|  | BOTW | HANZAB | BWD | Total | BOTW | HANZAB | Total |  |  |  |
| Passeriformes | 8 | 168 | 186 | 362 | 1348 | 191 | 1539 | 0.11 | 360 | 5.4 |
| Accipitriformes | 134 | 13 | 71 | 218 | 167 | 20 | 187 | 0.79 | 216 | 86.1 |
| Charadriiformes | 64 | 66 | 100 | 230 | 235 | 81 | 316 | 0.41 | 208 | 55.2 |
| Procellariiformes | 39 | 7 | 70 | 116 | 69 | 12 | 81 | 0.57 | 114 | 78.6 |
| Apodiformes | 1 | 7 | 80 | 88 | 68 | 7 | 75 | 0.11 | 88 | 18.4 |
| Anseriformes | 12 | 16 | 42 | 70 | 57 | 22 | 79 | 0.35 | 69 | 40.6 |
| Falconiformes | 33 | 5 | 19 | 57 | 38 | 5 | 43 | 0.88 | 54 | 84.4 |
| Psittaciformes | 1 | 45 | 2 | 48 | 273 | 51 | 324 | 0.14 | 48 | 11.9 |
| Pelecaniformes | 18 | 8 | 23 | 49 | 62 | 25 | 87 | 0.30 | 47 | 42.7 |
| Strigiformes | 7 | 7 | 26 | 40 | 142 | 7 | 149 | 0.09 | 40 | 16.8 |
| Suliformes | 13 | 9 | 17 | 39 | 26 | 9 | 35 | 0.63 | 36 | 67.9 |
| Gruiformes | 11 | 2 | 14 | 27 | 16 | 3 | 19 | 0.68 | 27 | 16.1 |
| Columbiformes | 0 | 17 | 9 | 26 | 142 | 36 | 178 | 0.10 | 24 | 6.8 |
| Coraciiformes | 6 | 11 | 4 | 21 | 143 | 15 | 158 | 0.11 | 21 | 11.2 |
| Galliformes<sup>†</sup> | 0 | 0 | 17 | 17 | 0 | 0 | 0 | - | 17 | 5.5 |
| Cuculiformes | 1 | 11 | 2 | 14 | 70 | 11 | 81 | 0.15 | 14 | 9.3 |
| Caprimulgiformes | 0 | 6 | 4 | 10 | 45 | 8 | 53 | 0.11 | 10 | 8.3 |
| Ciconiiformes | 5 | 0 | 5 | 10 | 15 | 0 | 15 | 0.33 | 10 | 50 |
| Piciformes | 0 | 0 | 8 | 8 | 168 | 0 | 168 | 0.00 | 8 | 1.7 |
| Otidiformes | 3 | 1 | 3 | 7 | 22 | 1 | 23 | 0.17 | 7 | 26.9 |
| Podicipediformes | 0 | 0 | 6 | 6 | 13 | 0 | 13 | 0.00 | 6 | 30 |
| Cathartiformes | 2 | 0 | 3 | 5 | 4 | 0 | 4 | 0.50 | 5 | 71.4 |
| Gaviiformes | 0 | 0 | 3 | 3 | 2 | 0 | 2 | 0.00 | 3 | 60 |
| Phaethontiformes | 0 | 0 | 3 | 3 | 0 | 0 | 0 | - | 3 | 100 |
| Phoenicopteriformes | 1 | 0 | 2 | 3 | 4 | 0 | 4 | 0.25 | 3 | 50 |
| Pterocliiformes | 0 | 0 | 2 | 2 | 0 | 0 | 0 | - | 2 | 12.5 |
| Bucerotiformes | 0 | 0 | 1 | 1 | 56 | 0 | 56 | 0.00 | 1 | 1.4 |
| Struthioniformes<sup>†</sup> | 0 | 0 | 1 | 1 | 1 | 0 | 1 | 0.00 | 1 | 1.6 |
| Coliiformes | 0 | 0 | 0 | 0 | 6 | 0 | 6 | 0.00 | 0 | 0 |
| Eurypygiformes | 0 | 0 | 0 | 0 | 2 | 0 | 2 | 0.00 | 0 | 0 |
| Leptosomiformes | 0 | 0 | 0 | 0 | 1 | 0 | 1 | 0.00 | 0 | 0 |
| Mesitornithiformes<sup>†</sup> | 0 | 0 | 0 | 0 | 3 | 0 | 3 | 0.00 | 0 | 0 |
| Musophagiformes | 0 | 0 | 0 | 0 | 24 | 0 | 24 | 0.00 | 0 | 0 |
| Opisthocomiformes | 0 | 0 | 0 | 0 | 1 | 0 | 1 | 0.00 | 0 | 0 |
| **Total** | **359** | **399** | **723** | **1481** | **3223** | **504** | **3727** | **0.20** | **1442** | **13.1** |

After removing flightless or poorly flighted orders, 1,424 species from 25 orders remained with wingspan data. Wing length was a strong predictor of wingspan across all species with empirical data (n = 1,424; R^2^ = 0.962, p < 0.001), with a global slope of 3.70 (intercept = −7.57), indicating that for every 1 cm increase in wing length, wingspan increases by approximately 3.70 cm. For the order-level models, we fitted 20 linear regressions, R^2^ values ranged from 0.705 (Caprimulgiformes, n = 10) to 0.992 (Piciformes, n = 8), with 17 of 20 order-level models achieving R^2^ ≥ 0.95 (Table 2). Slopes varied from 2.47 (Apodiformes) to 4.86 (Procellariiformes) (Fig. 1, Fig. 2; Appendix S2).

**Table 2:** Linear regression models predicting wingspan (cm) from wing length (cm) for 20 order-level groups with five or more species with empirical wingspan data, plus the global model fitted across all orders. Where an order had fewer than five species with empirical data, it was grouped with a functionally similar order prior to modelling (denoted by “&”). The global model was fitted across all 1,424 species with empirical wingspan data regardless of order. n = number of species used to fit the model; R^2^ = coefficient of determination. All models were statistically significant (p < 0.001). Predicted wingspans can be calculated by multiplying the slope by the target species folded wing measurement and adding the intercept as per a simple linear regression equation (y = mx + b).

| Order | n | Intercept | Slope | R<sup>2</sup> |
| --- | --- | --- | --- | --- |
| Piciformes | 8 | 1.65 | 3.014 | 0.992 |
| Gaviiformes & Podicipediformes | 9 | 10.25 | 3.692 | 0.989 |
| Apodiformes | 88 | -0.28 | 2.469 | 0.983 |
| Gruiformes | 27 | -2.85 | 3.722 | 0.978 |
| Coraciiformes | 21 | 1.75 | 3.296 | 0.975 |
| Procellariiformes | 114 | -34.03 | 4.864 | 0.971 |
| Strigiformes | 40 | -0.87 | 3.464 | 0.971 |
| Falconiformes | 54 | 0.00 | 3.047 | 0.970 |
| Pelecaniformes & Phoenicopteriformes | 50 | -12.83 | 4.248 | 0.967 |
| Accipitriformes | 221 | -14.76 | 3.712 | 0.965 |
| Passeriformes | 360 | 1.53 | 2.844 | 0.964 |
| Charadriiformes & Phaethontiformes | 211 | 1.68 | 3.220 | 0.958 |
| Otidiformes | 7 | 11.97 | 3.377 | 0.957 |
| Cuculiformes | 14 | 2.15 | 2.783 | 0.957 |
| Anseriformes | 69 | -6.77 | 3.840 | 0.955 |
| Columbiformes & Pterocliiformes | 26 | 0.65 | 2.989 | 0.951 |
| Psittaciformes | 48 | 1.64 | 2.858 | 0.949 |
| Suliformes | 36 | 22.41 | 3.180 | 0.944 |
| Ciconiiformes | 10 | -33.4 | 4.228 | 0.863 |
| Caprimulgiformes | 10 | 8.48 | 2.822 | 0.705 |
| **Global model** | **1424** | **-7.57** | **3.700** | **0.962** |

**Figure 1:**
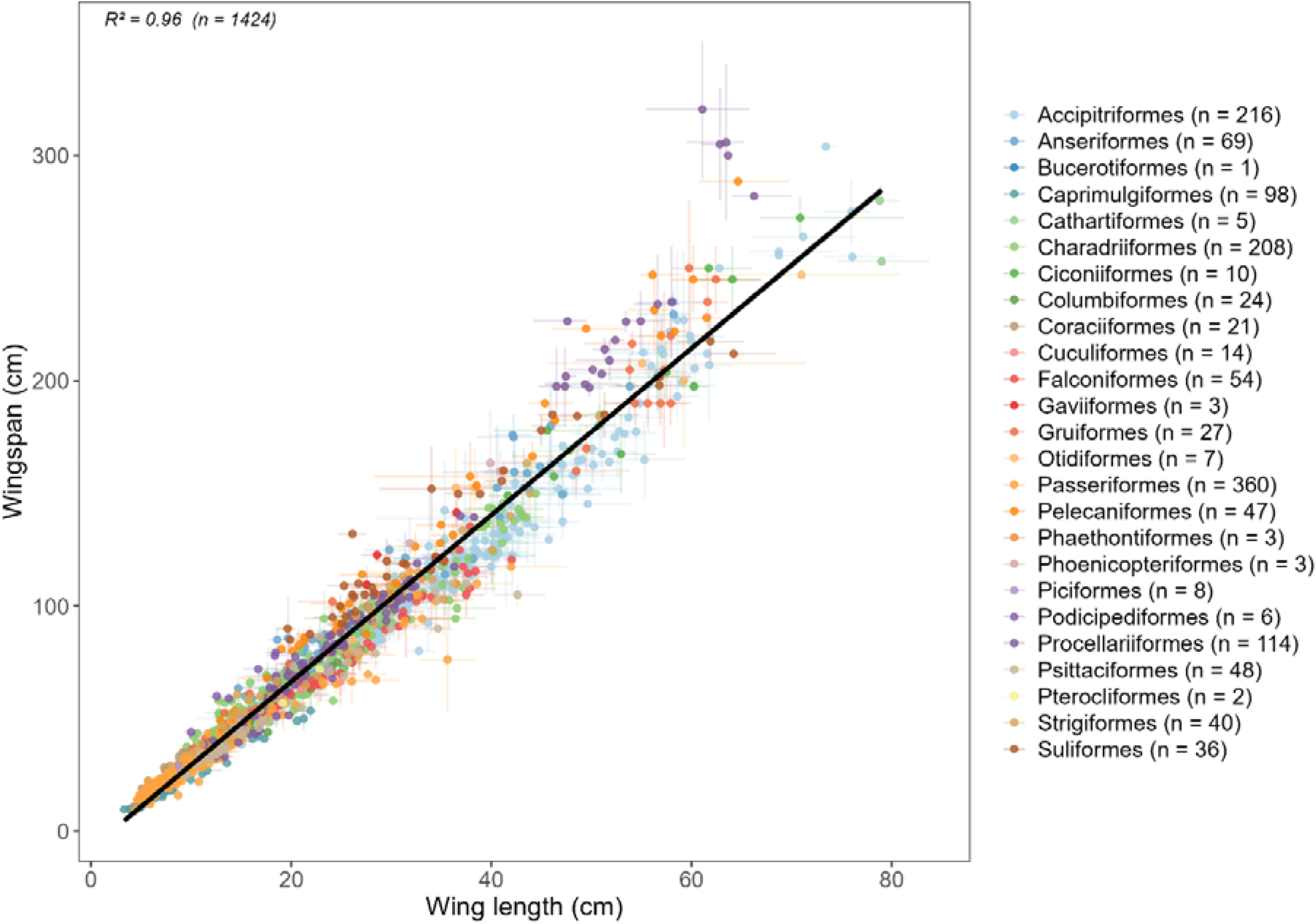
Relationship between wing length (cm) and wingspan (cm) across 1,424 species with empirical wingspan data, coloured by order. The black line represents the global linear regression fit (R^2^ = 0.96). Horizontal and vertical bars represent the range of wing length and wingspan measurements reported for each species, respectively.

**Figure 2:**
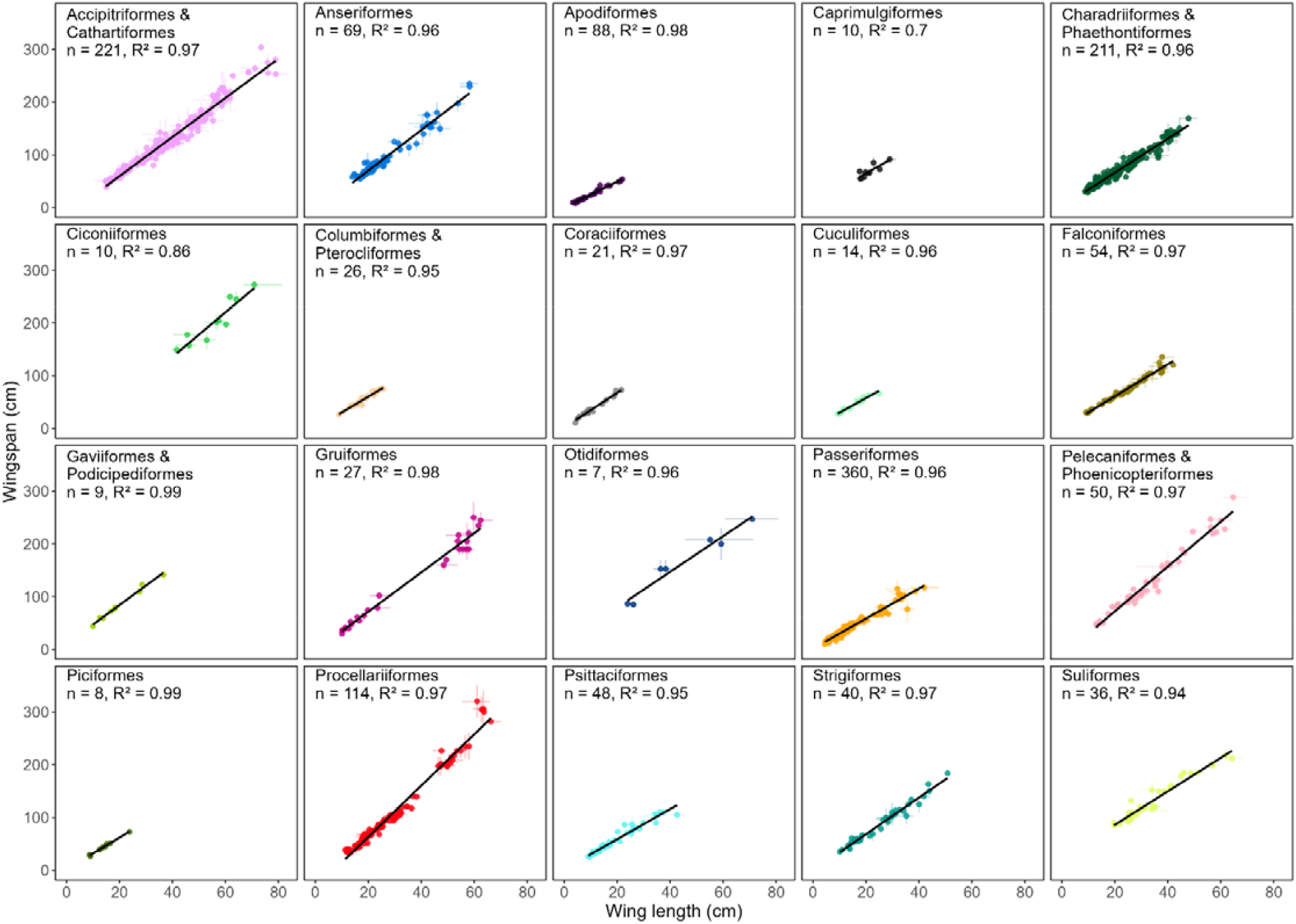
Order-level relationships between wing length (cm) and wingspan (cm), presented as individual panels with fixed axes across all panels to facilitate comparison of species size ranges among orders. The black line represents the fitted linear regression for each order. Horizontal and vertical bars represent the range of wing length and wingspan measurements reported for each species, respectively.

Across all order-level models, the median percentage deviation from CV was 4.9%, meaning that for a species with a predicted wingspan of 100 cm, the true wingspan would typically fall between 95.1 and 104.9 cm. Half of all predictions (50.7%) were within 5% of the empirical measured value, and 80% were within 10%. Across species with reported wingspan ranges (n = 941), the median intraspecific range (difference between midpoint and maximum or minimum value) was 5.7% (Appendix S3). For nine of 20 order-level models, model median deviation was less than or equal to the natural intraspecific variation, with an additional five order-level models having median deviation only slightly higher (0.5 percentage points) than intraspecific variation, suggesting that predictions are as accurate as the biological data would allow. Three orders had relatively higher model deviation compared to intraspecific variation, these were Caprimulgiformes (5.3 percentage point difference), Coraciiformes (3.7%), and Procellariiformes (3.2%) (Appendix S3).

Using the order-level and global models, wingspan was predicted for 9,194 flighted bird species lacking empirical data, producing a combined database of 10,618 species representing approximately 97% of all extant bird species. Order-level models were applied where available, with the global model used as a fallback for species belonging to orders without a fitted model (Bucerotiformes, Trogoniformes, Musophagiformes, Coliiformes, Eurypygiformes, Leptosomiformes, and Opisthocomiformes). The full predicted dataset, including predicted midpoint, minimum, and maximum wingspan values, is available in Appendix S1.

## DISCUSSION

Here we present a simple global method for predicting avian wingspan from wing length data, addressing a key data gap in collision risk modelling. As wing length data is available for all bird species (Tobias et al., 2022), we provide a reliable approach that can predict wingspans for flighted avian orders.

### Model performance and validation

Our cross-validation demonstrates that the order-level models presented here provide reliable wingspan estimates for flighted bird orders, with predictions falling close to the bounds of natural intraspecific variation for 14 of 20 orders. The global model and most order-level models indicate that wing length is a strong and consistent predictor of wingspan across avian diversity, consistent with the expected allometric scaling of wing dimensions (Nudds, 2007). The variation in slopes across orders reflects genuine biological differences in wing morphology, reinforcing the value of order-level rather than global-only modelling. Where species belong to orders not well represented in our dataset, the global model provides a reasonable fallback; however, users should consider whether the flight morphology of the focal species is likely to diverge substantially from the global mean. For example, species with long, narrow wings and high aspect ratios, such as dynamic-soaring seabirds or large soaring waterbirds, are likely to have wingspans underestimated by the global model, while compact-winged species may be overestimated.

Three orders exhibited relatively higher prediction deviation compared to intraspecific variation, warranting additional caution when applying model predictions. The Procellariiformes dataset represented 78.6% of species, and the order-level model achieved a high R^2^ (0.97); however, performance under cross-validation was poor.

Procellariiformes span an exceptional size gradient, from albatrosses (Diomedeidae) to small storm petrels (Hydrobatidae, Oceanitidae), encompassing a diverse array of flight strategies, from dynamic soaring of albatrosses to surface pattering of storm petrels (Spear & Ainley, 1997; Warham, 1977). The steep slope reflects the disproportionately long wings of Procellariiformes relative to other orders; the variation in wing architecture across the order means that a single regression line, while capturing the overall trend well, cannot fully account for the distinct wing length–wingspan relationships of each constituent family (see Appendix S2 for order-level figures grouped by family). Despite this, given the breadth of coverage and high R^2^, we consider the Procellariiformes model fit for purpose for generating wingspan estimates, with the caveat that family-level differences in wing morphology should be considered when interpreting predictions.

Caprimulgiformes had the poorest model fit (R^2^ = 0.705, n = 10) and was represented by two families, Caprimulgidae (nightjars) and Podargidae (frogmouths). Although these families are phylogenetically grouped within Caprimulgiformes, they differ in foraging behaviour and wing morphology: nightjars are aerial insectivores with long, pointed wings adapted for sustained aerial pursuit, while frogmouths are forest-dwelling sit-and-wait predators that hunt by pouncing on prey, with broader, more rounded wings (Winkler et al., 2024a, 2024b). These differences in wing shape likely drive the divergent wing length–wingspan ratios observed between families, producing greater scatter around the order-level regression line (Appendix S2). Given the poor fit and low coverage (10 of 120 species), we recommend using the global model predictions as a fallback for Caprimulgiformes until additional empirical wingspan data become available.

Coraciiformes achieved a high R^2^ (0.98) but had a relatively large difference between median prediction deviation and median intraspecific range. This discrepancy likely reflects the unusually low reported intraspecific variation for this order (median 3.4%) rather than poor model performance *per se*. The order encompasses families with distinct foraging strategies and wing morphologies, sit-and-wait plunge-diving kingfishers (Alcedinidae), acrobatic aerial-displaying rollers (Coraciidae), pursuit aerial insectivore bee-eaters (Meropidae), and the exceptionally small todies (Todidae, n = 1), suggesting that biological heterogeneity within the order contributes to residual scatter. We consider the Coraciiformes model broadly reliable but recommend caution for species belonging to Todidae, given their single-species representation in the training data.

Ciconiiformes achieved a moderate R^2^ (0.86) with only 10 species in the model data, spanning a wide size range from Abdim’s Stork (*Ciconia abdimii*; approximately 149 cm wingspan) to the Marabou Stork (*Leptoptilos crumenifer*; 263–282 cm wingspan). With few training data, the regression slope is particularly sensitive to the distribution of the available data, and predictions for poorly represented size classes may be unreliable. We recommend that predicted wingspans for Ciconiiformes are cross-checked against empirical measurements for morphologically similar species.

### Data coverage, geographic bias, and natural variation

Birds are one of the most comprehensively studied groups of vertebrates, encompassing enormous diversity in body size, flight strategy, foraging behaviour, and life history (Lees et al., 2022). Despite this research attention, wingspan data coverage was uneven across orders, indicating that wingspan is rarely measured or reported systematically outside of species where it has direct applied relevance. Our data collection efforts were prioritised towards orders identified as most at risk from wind turbine collisions, raptors, seabirds, and migratory waterbirds, and these orders are well represented in the final dataset (Accipitriformes 86.1%, Falconiformes 84.4%, Procellariiformes 78.6%). In contrast, several large and diverse orders remain poorly represented despite substantial search effort. Psittaciformes, the fourth largest avian order (Gill et al., 2025), yielded wingspan data for only 48 species (11.9%); similarly, Passeriformes, comprising over 60% of all bird species, is represented by only 5.4% of the order. Bucerotiformes, Musophagiformes, and Piciformes yielded no data other than a few species from *BirdWingData*. This uneven coverage reflects well-documented biases in avian research effort, which is skewed towards species of applied relevance, body size, and geographic range (Ducatez & Lefebvre, 2014; Titley et al., 2017). The bias in research effort is reflected in the availability of wingspan data; Psittaciformes, Columbiformes, and Piciformes are among the most understudied avian orders relative to their species richness, and their diversity is concentrated in tropical regions that are systematically understudied in biodiversity research (Titley et al., 2017). Consequently, data availability is poor for species in these regions, highlighting the importance of our approach to predicting wingspan for species without empirical data.

As expected, order-level slopes varied considerably, reflecting genuine differences in wing morphology across avian diversity. Wingspan, like other morphological traits, varies naturally within species due to intraspecific factors including geographic variation along climatic gradients (Bergmann’s rule; Meiri & Dayan, 2003) and sexual dimorphism (Caron & Pie, 2025). Any predicted wingspan therefore represents a point estimate within a broader biological range, and users should treat predictions accordingly — particularly for species with pronounced sexual size dimorphism or wide geographic distributions.

### Applications

The wingspan estimates generated here have immediate practical value for collision risk modelling, where wingspan is a required input for widely used models (e.g., Band & Band, 2012; Band et al., 2007; Smales et al., 2013), and is used to estimate flight speed (KleinHeerenbrink et al., 2015; Pennycuick, 2008). For practitioners conducting environmental impact assessments, these models provide the first validated method for obtaining wingspan estimates for species lacking empirical data, supporting more quantitative and species-specific risk assessments. This is particularly valuable for expanding collision risk assessments into the Southern Hemisphere where wind energy development is accelerating, and morphological data for Procellariiformes, Psittaciformes, and other relevant taxa are often sparse (Dunnett et al., 2022; Miller et al., 2025; Reid et al., 2025). Beyond collision risk, wingspan is a key determinant of flight performance relevant to migration modelling, flight energetics (Hedenström, 2025), and macroecological analyses of wing morphology and its ecological correlates.

## CONCLUSION

Here we present order-level linear regression models for predicting avian wingspan from wing length, validated through cross-validation and contextualised against intraspecific variation in wingspan. Wing length is available for all extant bird species through AVONET (Tobias et al., 2022), and we have used these models to generate wingspan estimates for all flighted bird species, effectively closing a key data gap in avian morphology. The models perform well across orders, with predictions typically falling within the bounds of intraspecific variation and perform well for the orders of greatest relevance in the context of wind energy development — raptors, seabirds, and migratory waterbirds. For practitioners conducting collision risk assessments, these models provide a transparent and reproducible method for obtaining wingspan estimates where empirical data are unavailable, supporting more quantitative and species-specific risk assessments. More broadly, the accompanying wingspan dataset represents the most comprehensive compilation of empirical and predicted avian wingspan data to date, and together with the regression models, serves as useful resources for macroecological analyses of flight morphology and its ecological correlates. As wind energy development continues to expand globally, tools that extend morphological data coverage to data-poor species will become increasingly important for balancing renewable energy expansion with biodiversity conservation.

## Supplementary materials

**Appendix S1:** Empirical and predicted wingspan xlsx dataset.

**Appendix S2:** Individual order-level figures colour-coded by family.

**Appendix S3:** Table S1 and Figure S1.

## Data Availability

The data and code that support the findings of this study are openly available from GitHub at https://github.com/MichaelRihaFox/project_wingspan. The compiled wingspan database is available as Appendix S1 and deposited on FigShare at 10.6084/m9.figshare.32089083.

## References

Alerstam, T., Rosén, M., Bäckman, J., Ericson, P. G. P., & Hellgren, O. (2007). Flight Speeds among Bird Species: Allometric and Phylogenetic Effects. PLOS Biology, 5(8), e197. 10.1371/journal.pbio.0050197.

Band, B., & Band, B. (2012). Using a collision risk model to assess bird collision risks for offshore windfarms. British Trust for Ornithology.

Band, W., Madders, M., & Whitfield, P. D. (2007). Developing field and analytical methods to assess avian collision risk at wind farms. In M. de Lucas, G. F. E. Janss, & M. Ferrer (Eds.), Birds and wind farms: risk assessment and mitigation (pp. 259–275).

Beston, J. A., Diffendorfer, J. E., Loss, S. R., & Johnson, D. H. (2016). Prioritizing Avian Species for Their Risk of Population-Level Consequences from Wind Energy Development. PLOS ONE, 11(3), e0150813. doi:10.1371/journal.pone.0150813.

Biasotto, L. D., Moreira, F., Bencke, G. A., D’Amico, M., Kindel, A., & Ascensão, F. (2022). Risk of bird electrocution in power lines: a framework for prioritizing species and areas for conservation and impact mitigation. Animal Conservation, 25(2), 285–296. 10.1111/acv.12736.

BirdLife Australia. (2023). 1990-2006 Handbook of Australian, New Zealand and Antarctic Birds: Volume 1 to 7. In S. Marchant & P. J. Higgins (Eds.). https://hanzab.birdlife.org.au/

BirdLife International. (2025). AVISTEP: the Avian Sensitivity Tool for Energy Planning. Technical Manual. B. International.

Caron, F. S., & Pie, M. R. (2025). The macroevolution of sexual size dimorphism in birds. Biological Journal of the Linnean Society, 144(3), blad168. 10.1093/biolinnean/blad168.

Cook, A. S. C. P., Salkanovic, E., Masden, E., Lee, H. E., & Kiilerich, A. H. (2025). A critical appraisal of 40 years of avian collision risk modelling: How have we got here and where do we go next? Environmental Impact Assessment Review, 110, 107717.

Cornell Lab of Ornithology. (2026). Birds of the World (M. Billerman, Ed.) https://birdsoftheworld.org/bow/home

Desholm, M. (2009). Avian sensitivity to mortality: Prioritising migratory bird species for assessment at proposed wind farms. Journal of Environmental Management, 90(8), 2672–2679. 10.1016/j.jenvman.2009.02.005.

Ducatez, S., & Lefebvre, L. (2014). Patterns of Research Effort in Birds. PLOS ONE, 9(2), e89955. doi:10.1371/journal.pone.0089955.

Dunnett, S., Holland, R. A., Taylor, G., & Eigenbrod, F. (2022). Predicted wind and solar energy expansion has minimal overlap with multiple conservation priorities across global regions. Proceedings of the National Academy of Sciences, 119(6), e2104764119. 10.1073/pnas.2104764119.

Estellés-Domingo, I., & López-López, P. (2025). Effects of wind farms on raptors: A systematic review of the current knowledge and the potential solutions to mitigate negative impacts. Animal Conservation, 28(3), 334–352. 10.1111/acv.12988.

Fu, H., Su, M., Chu, J. J., Margaritescu, A., & Claramunt, S. (2023). New methods for estimating the total wing area of birds. Ecology and Evolution, 13(9), e10480. 10.1002/ece3.10480.

Furness, R. W., Wade, H. M., & Masden, E. A. (2013). Assessing vulnerability of marine bird populations to offshore wind farms. Journal of Environmental Management, 119, 56–66. 10.1016/j.jenvman.2013.01.025.

Gémard, C., Duriez, O., Chappe, O., Duclos, G., & Besnard, A. (2025). Towards a better understanding of avian collision in wind energy facilities using automatic detection systems. Journal of Applied Ecology, 62(6), 1437–1448. 10.1111/1365-2664.70055.

Gill, F., Donsker, D., & Rasmussen, P. (2025). IOC World Bird List (v15.1) (https://www.worldbirdnames.org/

Grilli, M. G., Lambertucci, S. A., Therrien, J.-F., & Bildstein, K. L. (2017). Wing size but not wing shape is related to migratory behavior in a soaring bird. Journal of Avian Biology, 48(5), 669–678. 10.1111/jav.01220.

Hedenström, A. (2025). Integrating flight mechanics, energetics and migration ecology in vertebrates. Journal of Experimental Biology, 228(Suppl_1). doi:10.1242/jeb.248123.

Klein Heerenbrink, M. (2023). afpt: Tools for modelling of animal flight performance. R package (Version 1.0.3). 10.32614/CRAN.package.afpt.

KleinHeerenbrink, M., Johansson, L. C., & Hedenström, A. (2015). Power of the wingbeat: modelling the effects of flapping wings in vertebrate flight. Proceedings of the Royal Society A: Mathematical, Physical and Engineering Sciences, 471(2177), 20140952. doi:10.1098/rspa.2014.0952.

Lees, A. C., Haskell, L., Allinson, T., Bezeng, S. B., Burfield, I. J., Renjifo, L. M., Rosenberg, K. V., Viswanathan, A., & Butchart, S. H. M. (2022). State of the World’s Birds. Annual Review of Environment and Resources, 47, 231–260. 10.1146/annurev-environ-112420-014642.

Masden, E. A., & Cook, A. S. C. P. (2016). Avian collision risk models for wind energy impact assessments. Environmental Impact Assessment Review, 56, 43–49. 10.1016/j.eiar.2015.09.001.

Masden, E. A., Cook, A. S. C. P., McCluskie, A., Bouten, W., Burton, N. H. K., & Thaxter, C. B. (2021). When speed matters: The importance of flight speed in an avian collision risk model. Environmental Impact Assessment Review, 90, 106622. 10.1016/j.eiar.2021.106622.

Meiri, S., & Dayan, T. (2003). On the validity of Bergmann’s rule. Journal of Biogeography, 30(3), 331–351. 10.1046/j.1365-2699.2003.00837.x.

Miller, M. G. R., Petrovic, S., & Clarke, R. H. (2025). A global review of Procellariiform flight height, flight speed and nocturnal activity: Implications for offshore wind farm collision risk. Journal of Applied Ecology, 62(8), 1795–1819. 10.1111/1365-2664.70088.

Nudds, R. L. (2007). Wing-bone length allometry in birds. Journal of Avian Biology, 38(4), 515–519. 10.1111/j.0908-8857.2007.03913.x.

Pennycuick, C. J. (2008). Modelling the flying bird (Vol. 5). Elsevier.

R Core Team. (2025). R: A Language and Environment for Statistical Computing. R Foundation for Statistical Computing. https://www.R-project.org/.

Reid, K., Baker, G. B., & Ehmke, G. (2025). Assessing impacts on birds from onshore wind farms in Australia: an ecological risk assessment. Emu - Austral Ornithology, 1–15. 10.1080/01584197.2025.2582504.

Reid, K., Baker, G. B., & Woehler, E. J. (2023). An ecological risk assessment for the impacts of offshore wind farms on birds in Australia. Austral Ecology, 48(2), 418–439. 10.1111/aec.13278.

Sheard, C., Neate-Clegg, M. H. C., Alioravainen, N., Jones, S. E. I., Vincent, C., MacGregor, H. E. A., Bregman, T. P., Claramunt, S., & Tobias, J. A. (2020). Ecological drivers of global gradients in avian dispersal inferred from wing morphology. Nature communications, 11(1), 2463. 10.1038/s41467-020-16313-6.

Shiomi, K., Tatani, M., & Kikuchi, D. M. (2025). BirdWingData: Wingspan and wing area data of birds compiled from multiple literature sources and original measurements. Ecological Research, 40(1), 82–89. 10.1111/1440-1703.12502.

Smales, I., Muir, S., Meredith, C., & Baird, R. (2013). A description of the biosis model to assess risk of bird collisions with wind turbines. Wildlife Society Bulletin, 37(1), 59–65. 10.1002/wsb.257.

Spear, L. B., & Ainley, D. G. (1997). Flight behaviour of seabirds in relation to wind direction and wing morphology. Ibis, 139(2), 221–233. 10.1111/j.1474-919X.1997.tb04620.x.

Sullivan, T. N., Meyers, M. A., & Arzt, E. (2019). Scaling of bird wings and feathers for efficient flight. Science Advances, 5(1), eaat4269. 10.1126/sciadv.aat4269.

Thaxter, C. B., Buchanan, G. M., Carr, J., Butchart, S. H., Newbold, T., Green, R. E., Tobias, J. A., Foden, W. B., O’Brien, S., & Pearce-Higgins, J. W. (2017). Bird and bat species’ global vulnerability to collision mortality at wind farms revealed through a trait-based assessment. Proceedings of the Royal Society B: Biological Sciences, 284(1862), 20170829. https://pmc.ncbi.nlm.nih.gov/articles/PMC5597824/

Titley, M. A., Snaddon, J. L., & Turner, E. C. (2017). Scientific research on animal biodiversity is systematically biased towards vertebrates and temperate regions. PLOS ONE, 12(12), e0189577. doi:10.1371/journal.pone.0189577.

Tobias, J. A., Sheard, C., Pigot, A. L., Devenish, A. J. M., Yang, J., Sayol, F., Neate-Clegg, M. H. C., Alioravainen, N., Weeks, T. L., Barber, R. A., Walkden, P. A., MacGregor, H. E. A., Jones, S. E. I., Vincent, C., Phillips, A. G., Marples, N. M., Montaño-Centellas, F. A., Leandro-Silva, V., Claramunt, S.,…Schleuning, M. (2022). AVONET: morphological, ecological and geographical data for all birds. Ecology Letters, 25(3), 581–597. 10.1111/ele.13898.

Warham, J. (1977). Wing loadings, wing shapes, and flight capabilities of procellariiformes. New Zealand Journal of Zoology, 4(1), 73–83. 10.1080/03014223.1977.9517938.

Winkler, D. W., Billerman, S. M., & Lovette, I. J. (2024a). Frogmouths (Podargidae), version 2.0. In S. M. Billerman, B. K. Keeney, P. G. Rodewald, & T. S. Schulenberg (Eds.), Birds of the World. Cornell Lab of Ornithology. 10.2173/bow.podarg1.02.

Winkler, D. W., Billerman, S. M., & Lovette, I. J. (2024b). Nightjars and Allies (Caprimulgidae), version 2.0. In S. M. Billerman, B. K. Keeney, P. G. Rodewald, & T. S. Schulenberg (Eds.), Birds of the World. Cornell Lab of Ornithology. 10.2173/bow.caprim2.02.

