## Appendix S2 for "Estimating avian wingspan from wing length: an order-level regression approach"

**Appendix S2**: Order-level relationships between wing length (cm) and wingspan (cm) for each of the 20 order-level model groups. Points are coloured by family. The black line represents the fitted linear regression for each order group. Horizontal and vertical bars represent the range of wing length and wingspan measurements reported for each species, respectively. Where orders with insufficient data were grouped with a functionally or phylogenetically similar order, both order names are shown in the figure title. Model fit (R²) and sample size (n) are shown in each panel title.

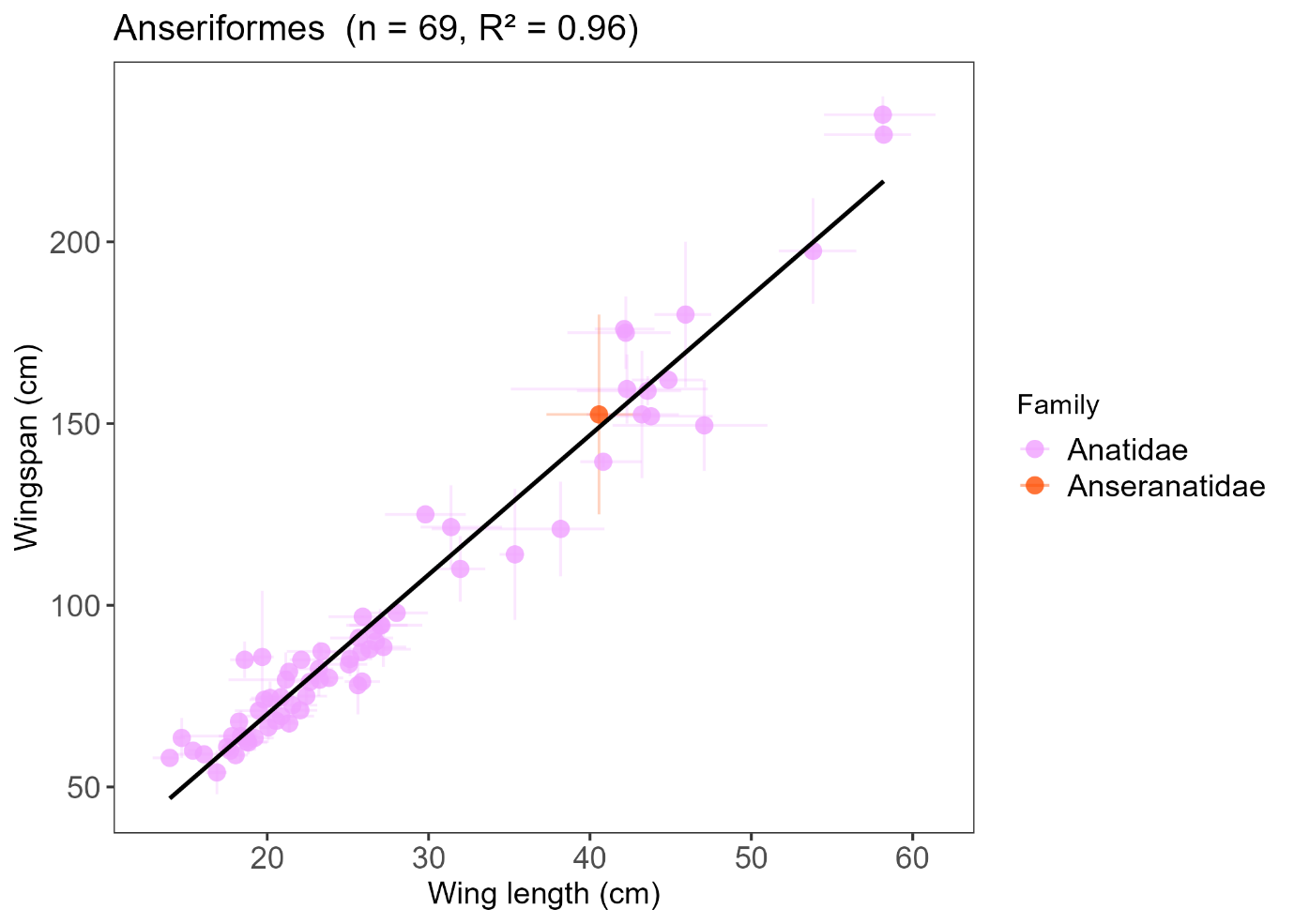

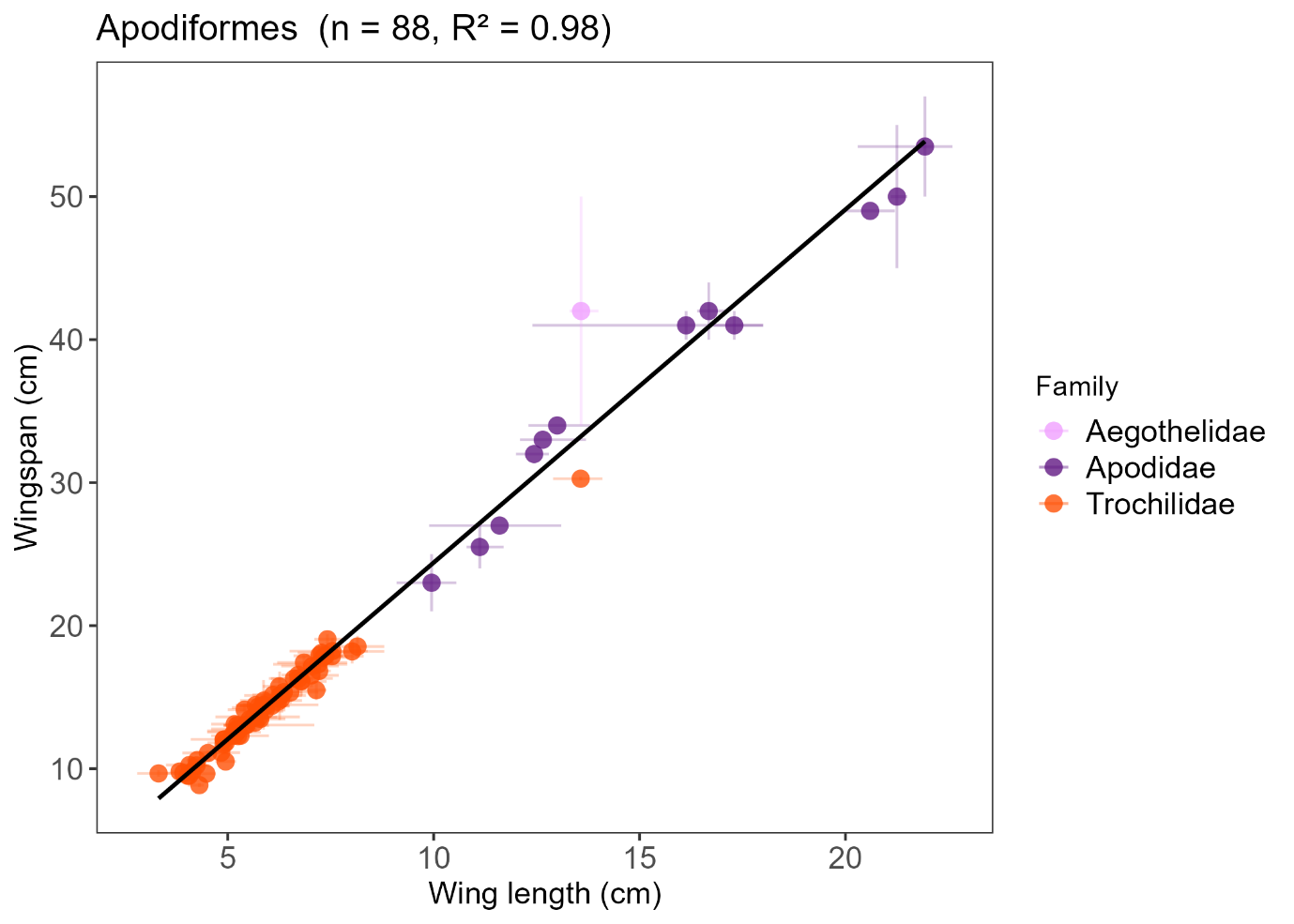

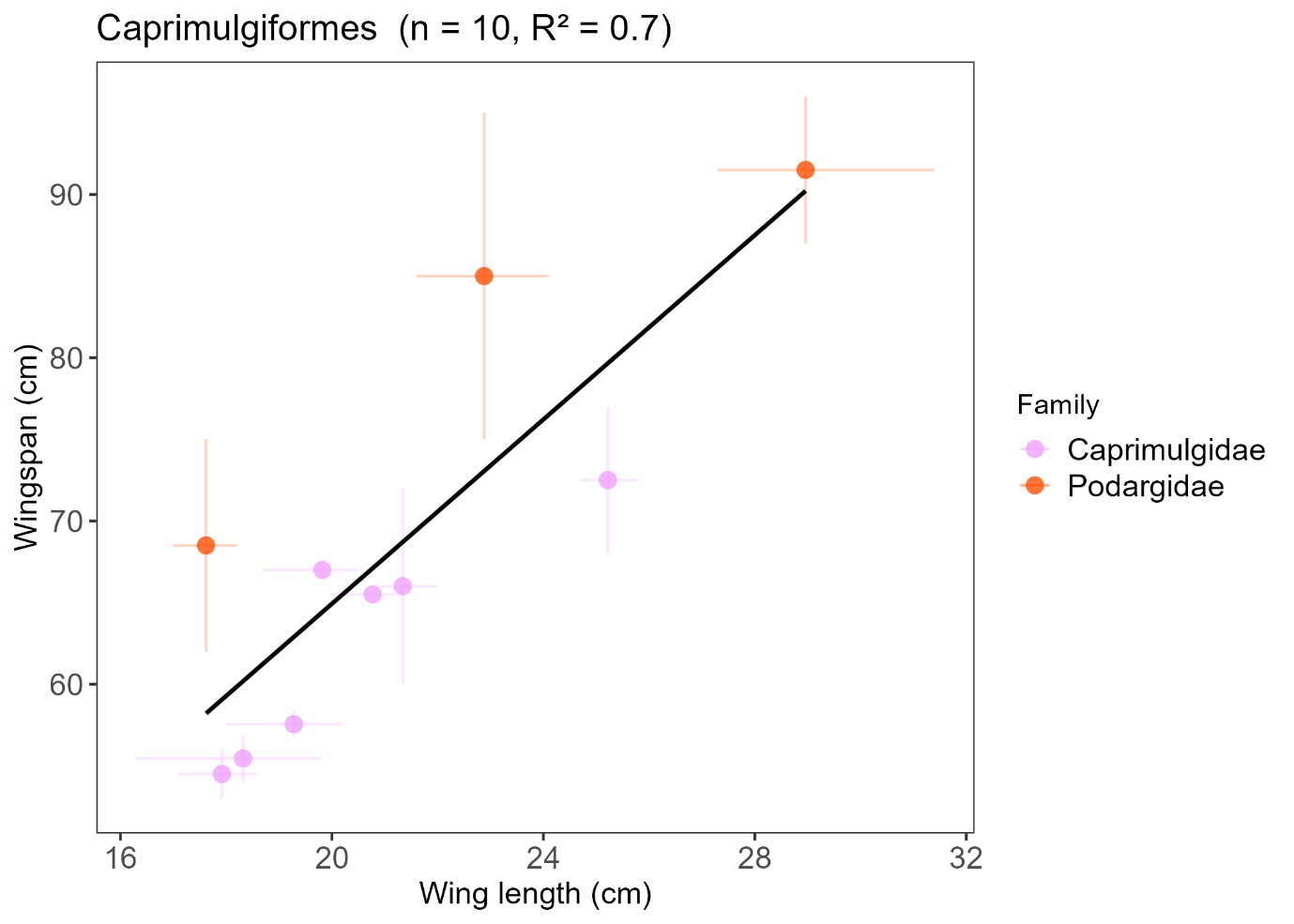

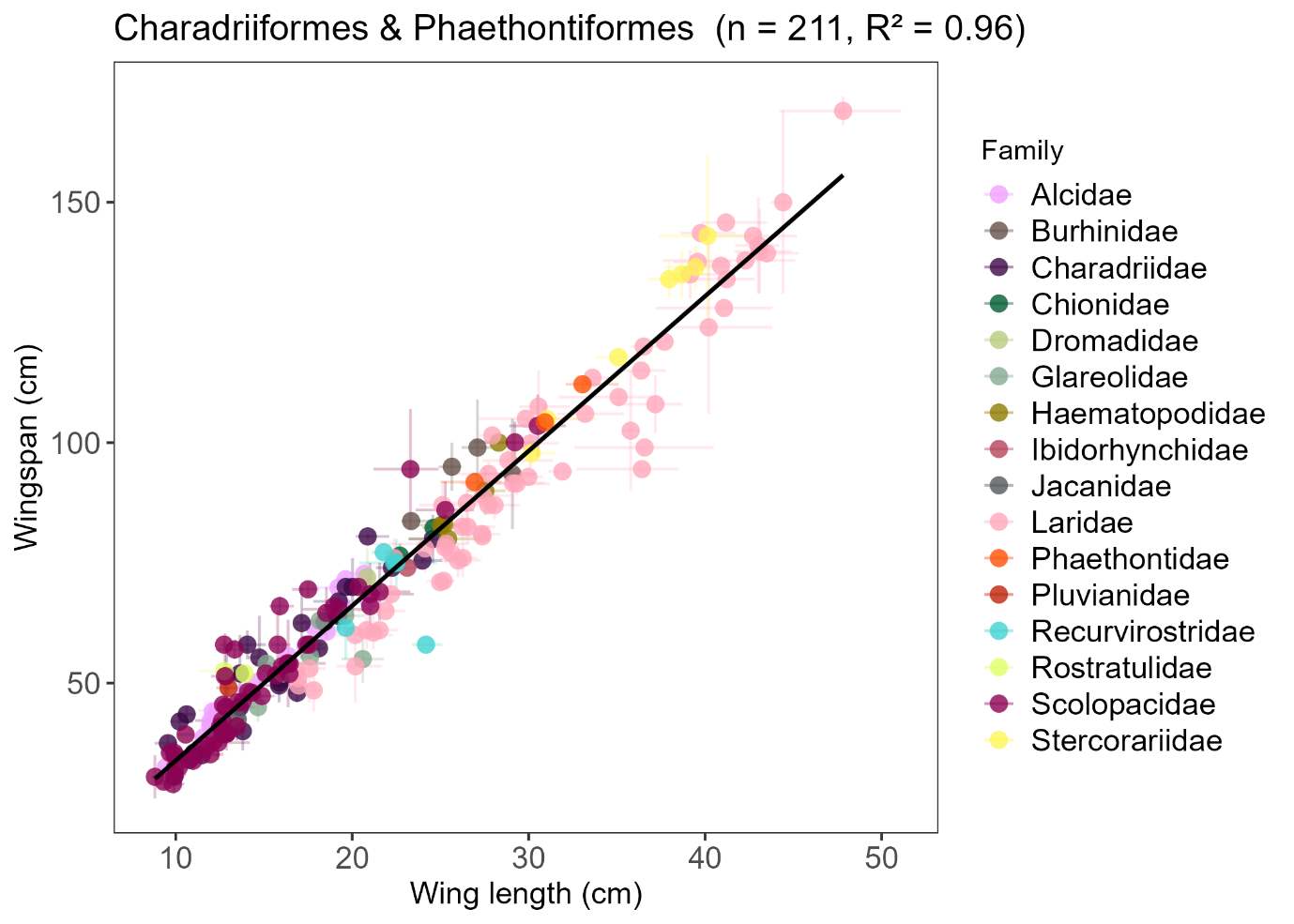

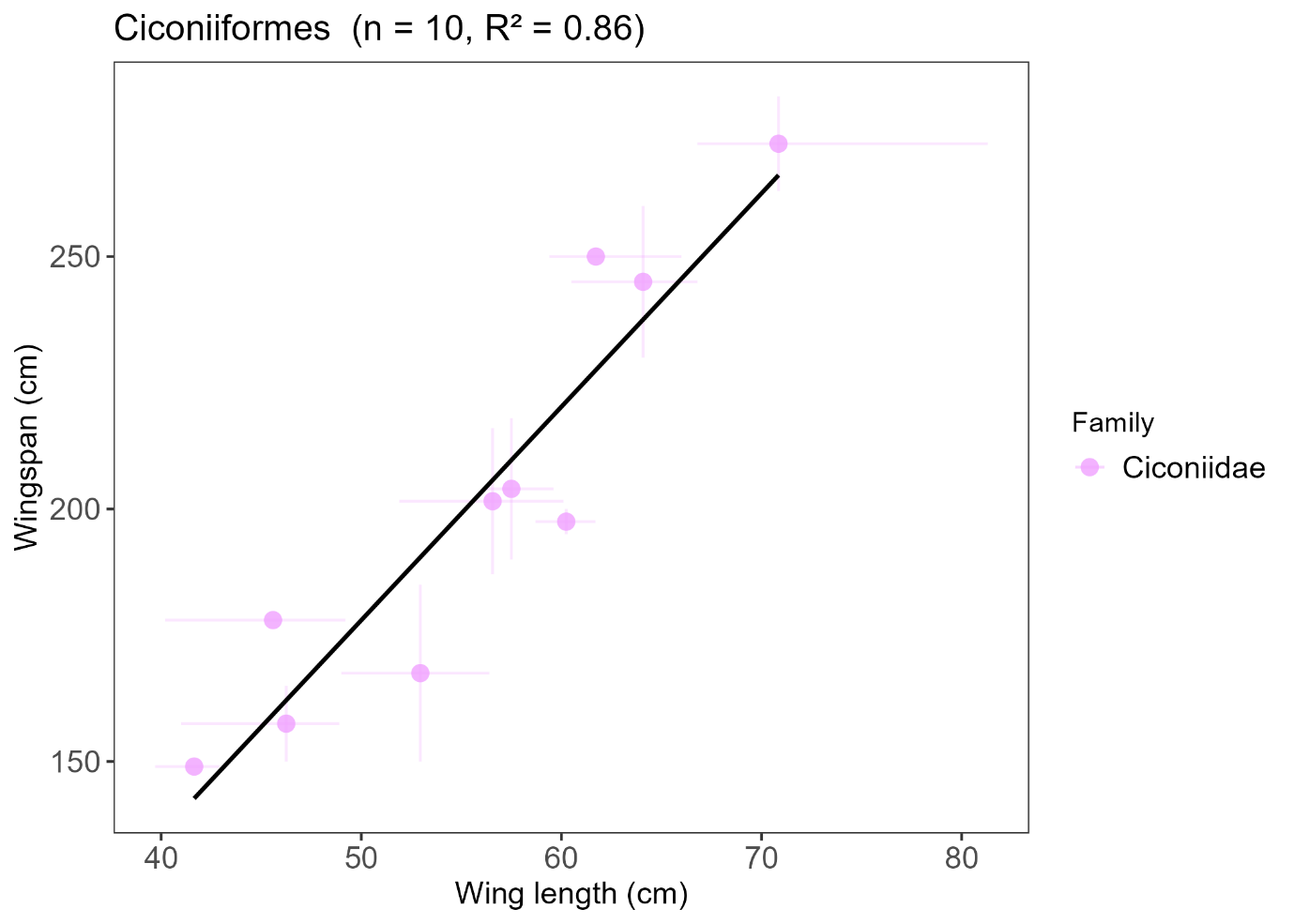

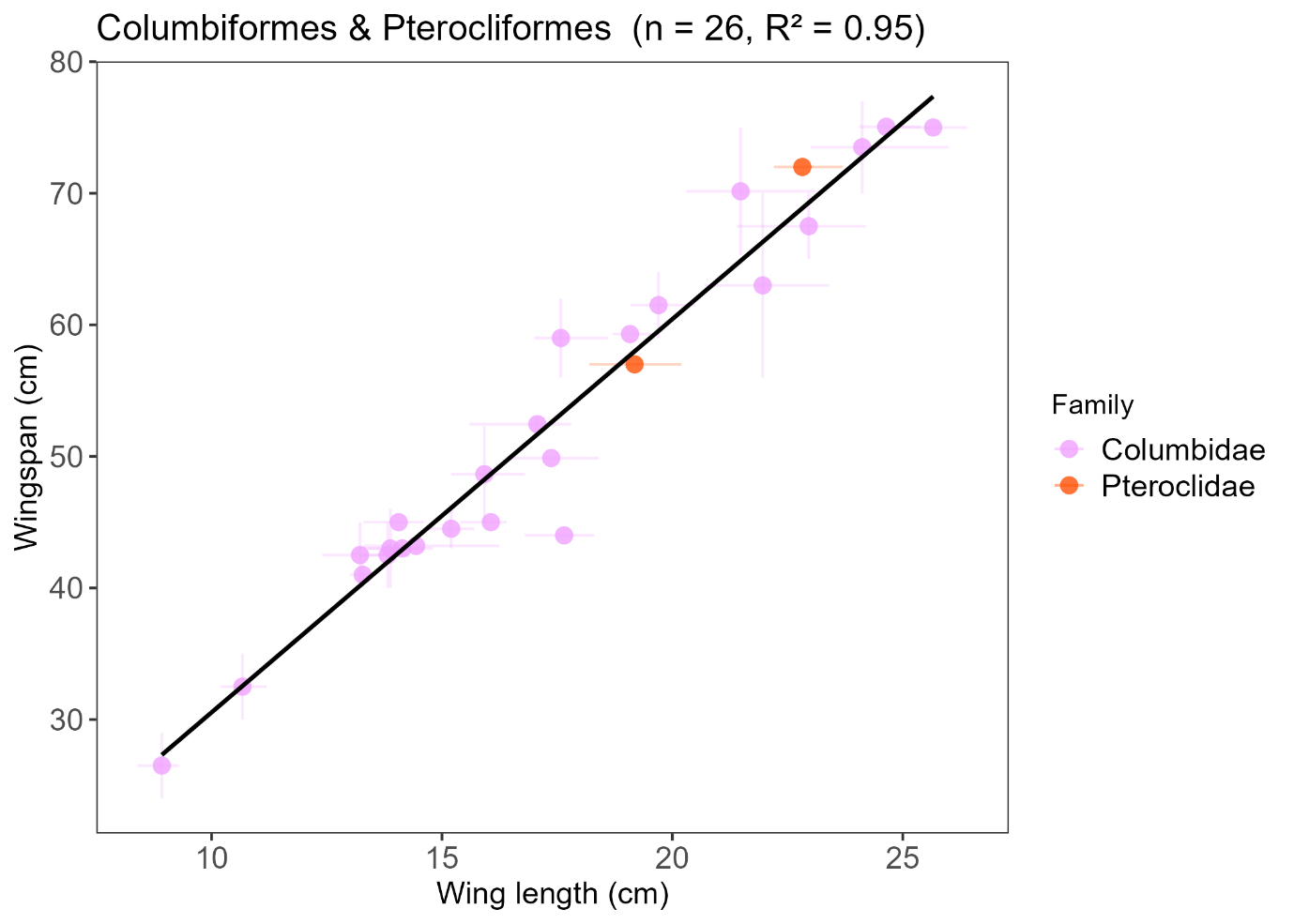

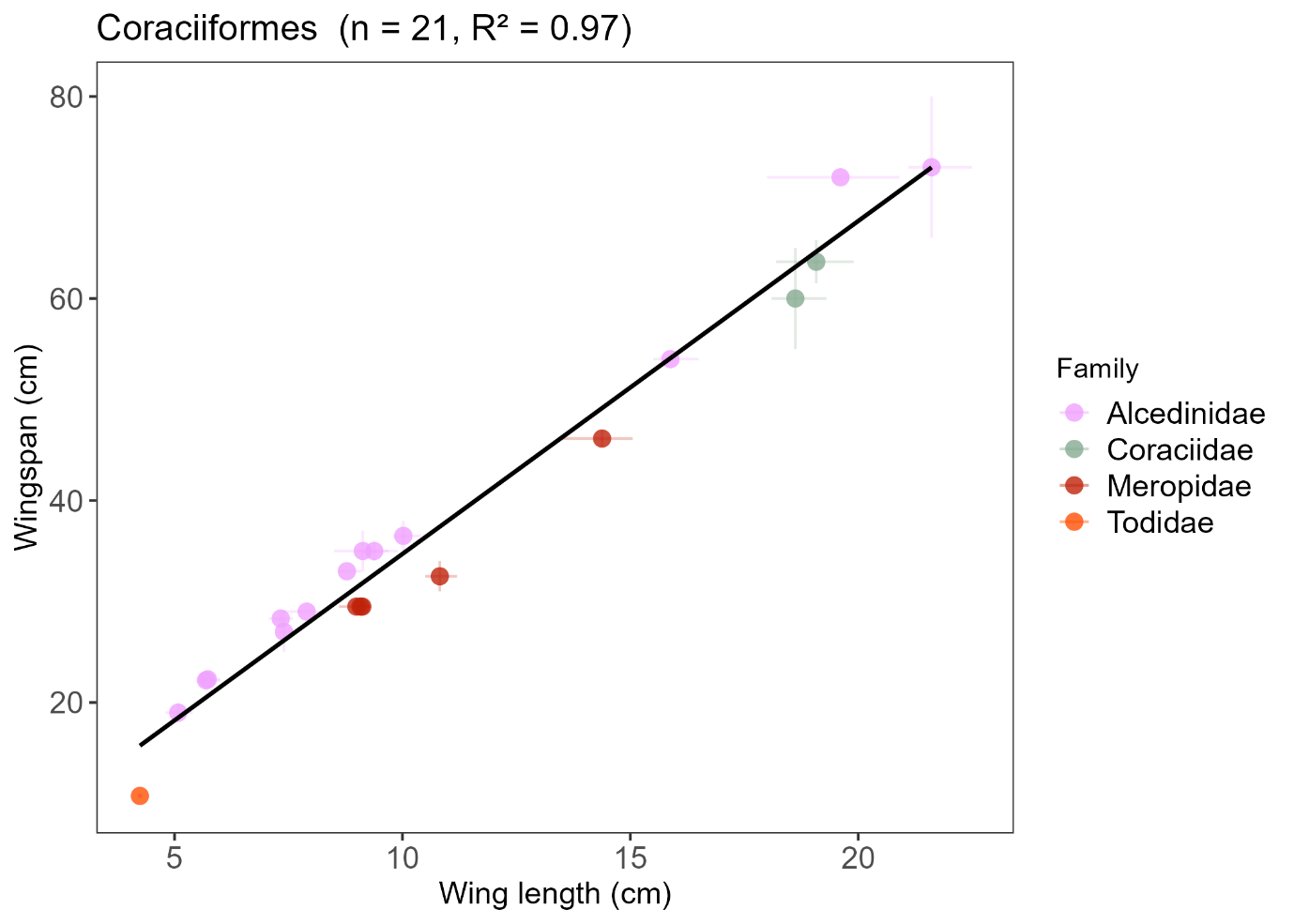

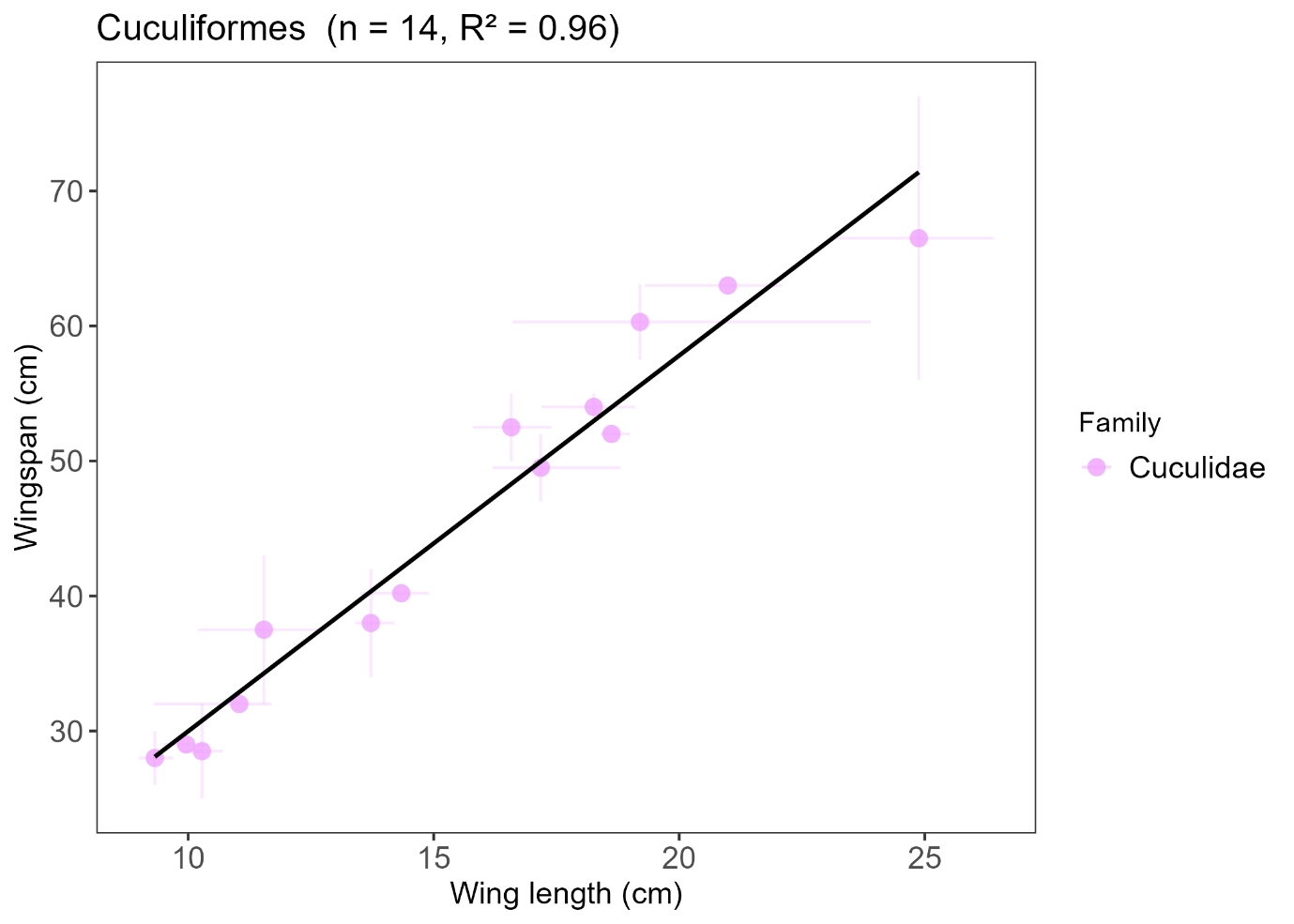

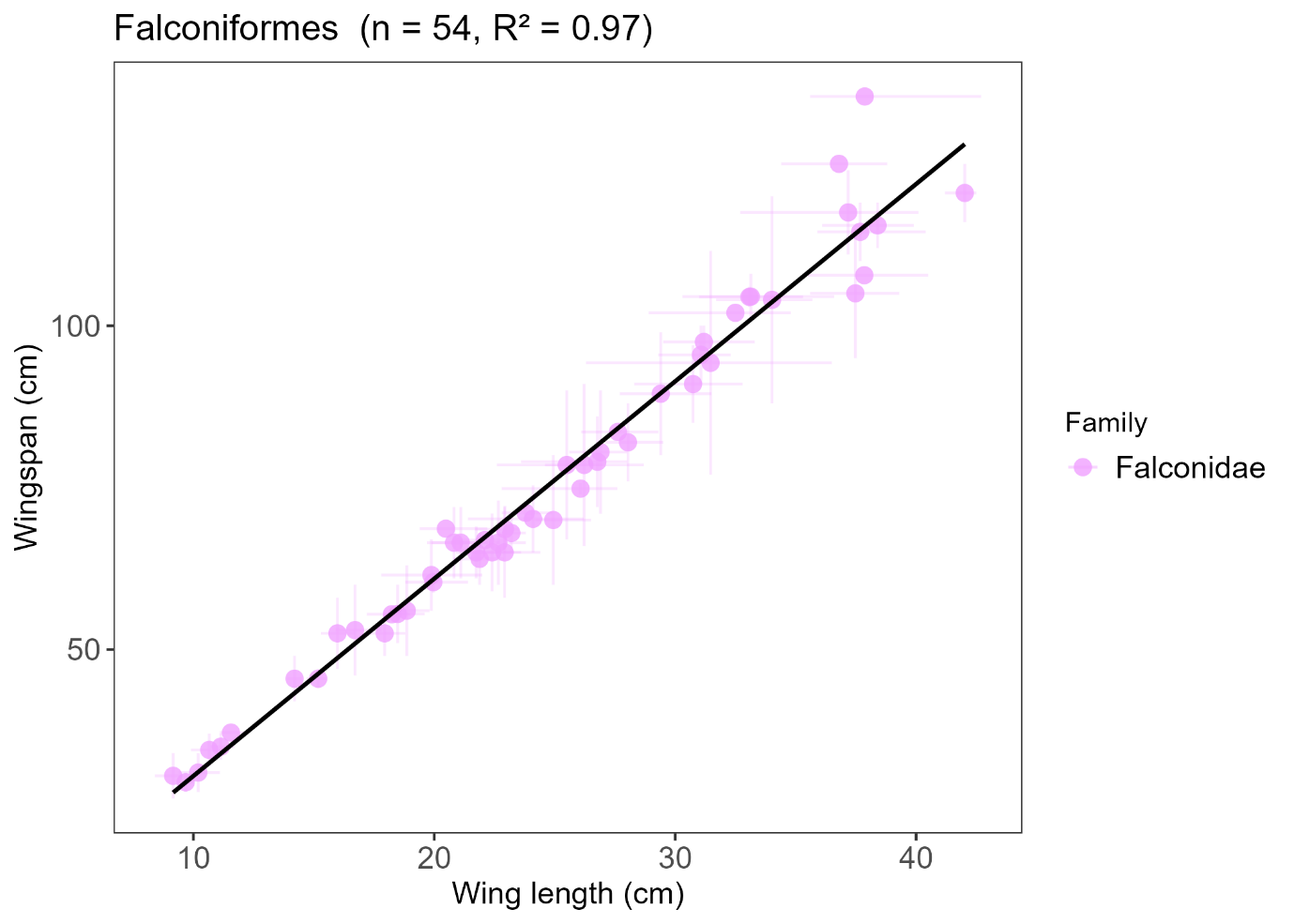

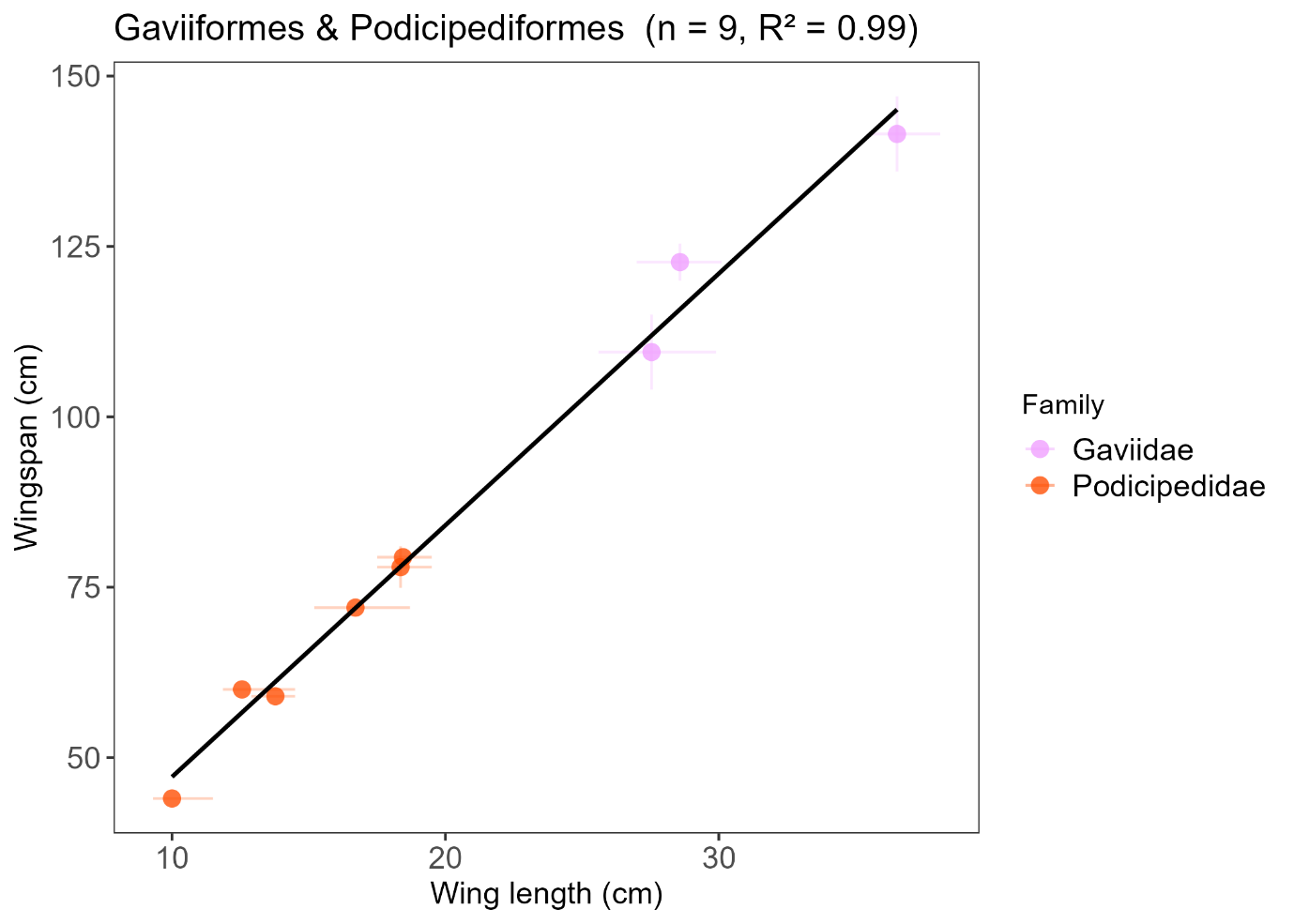

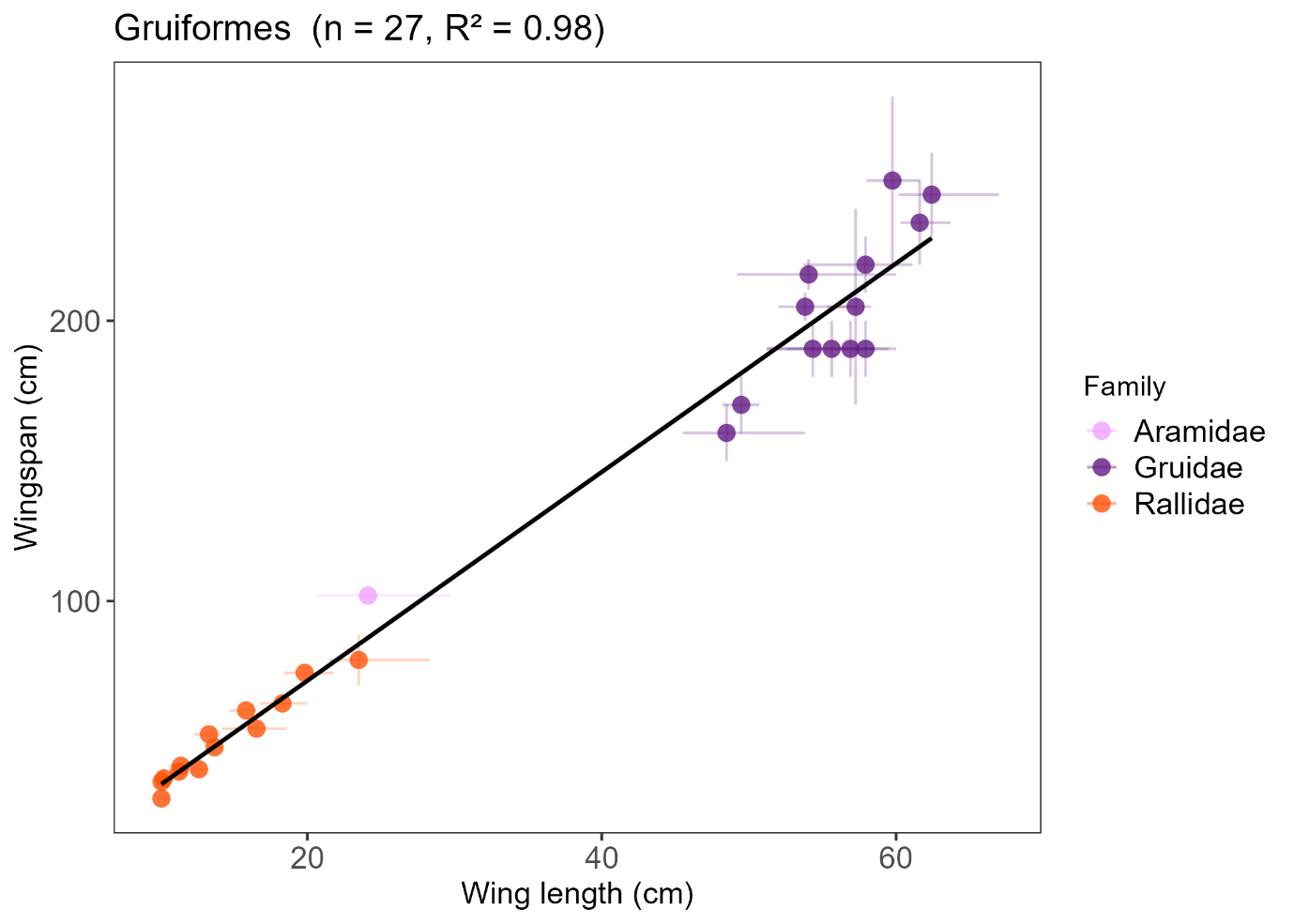

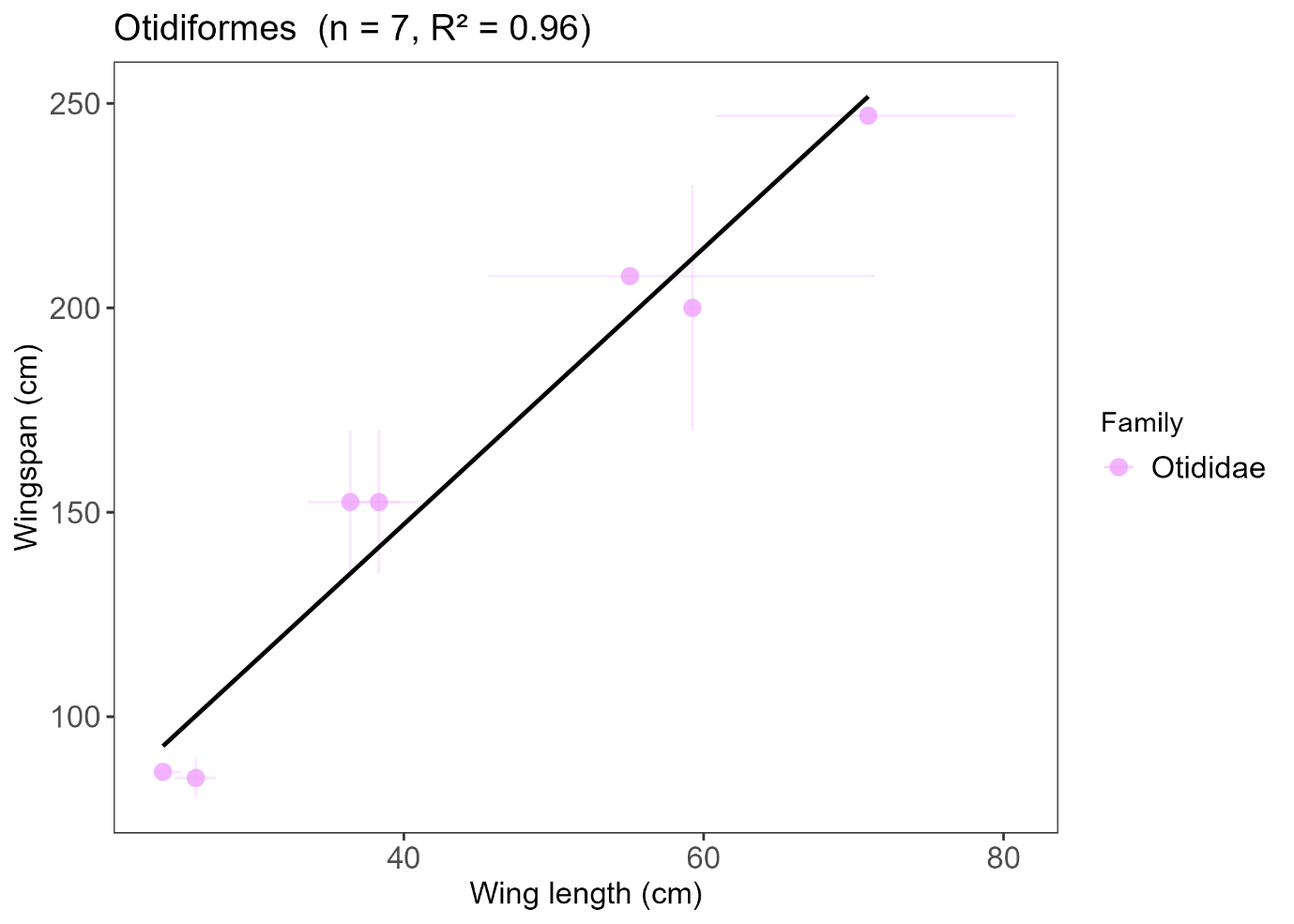

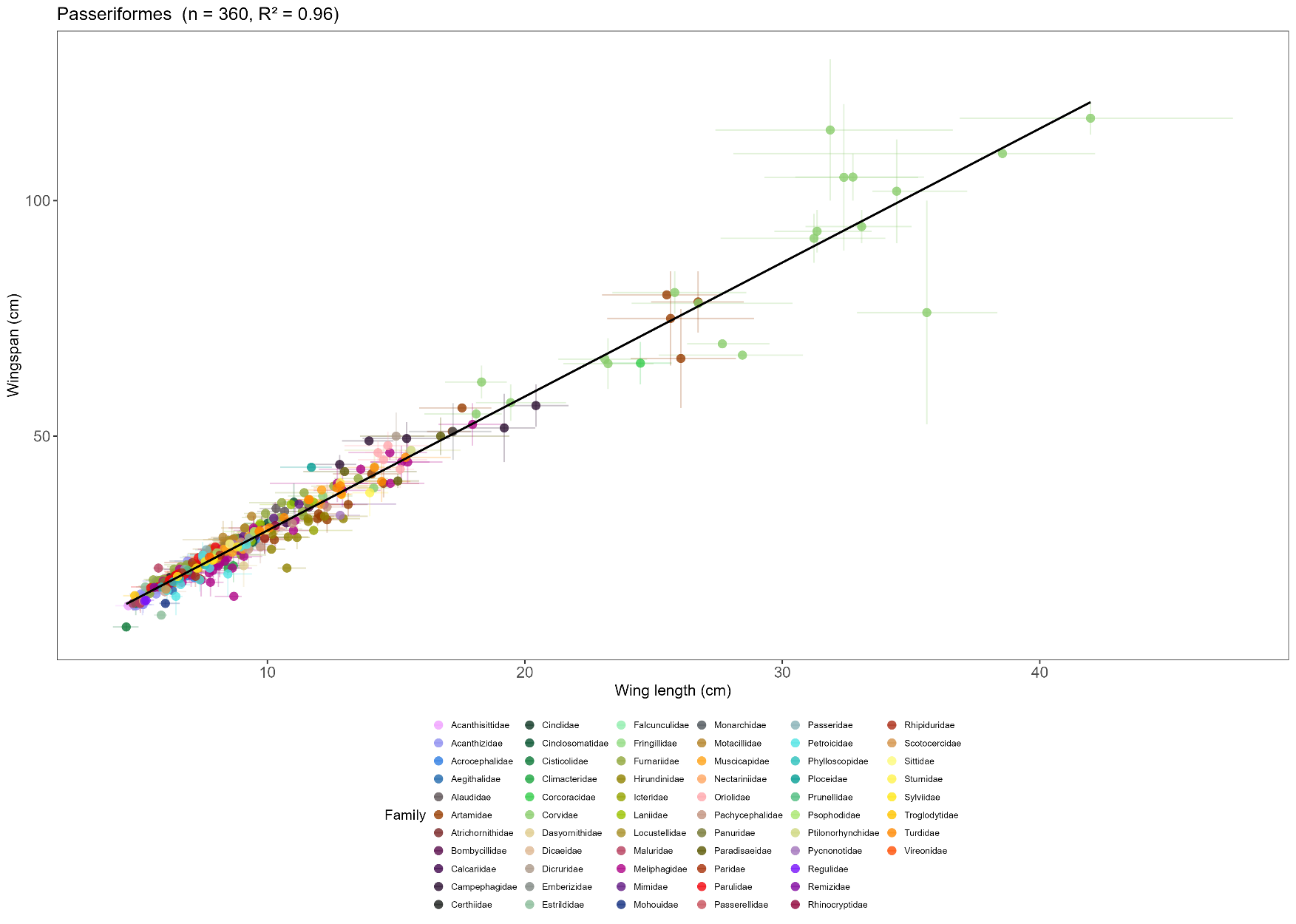

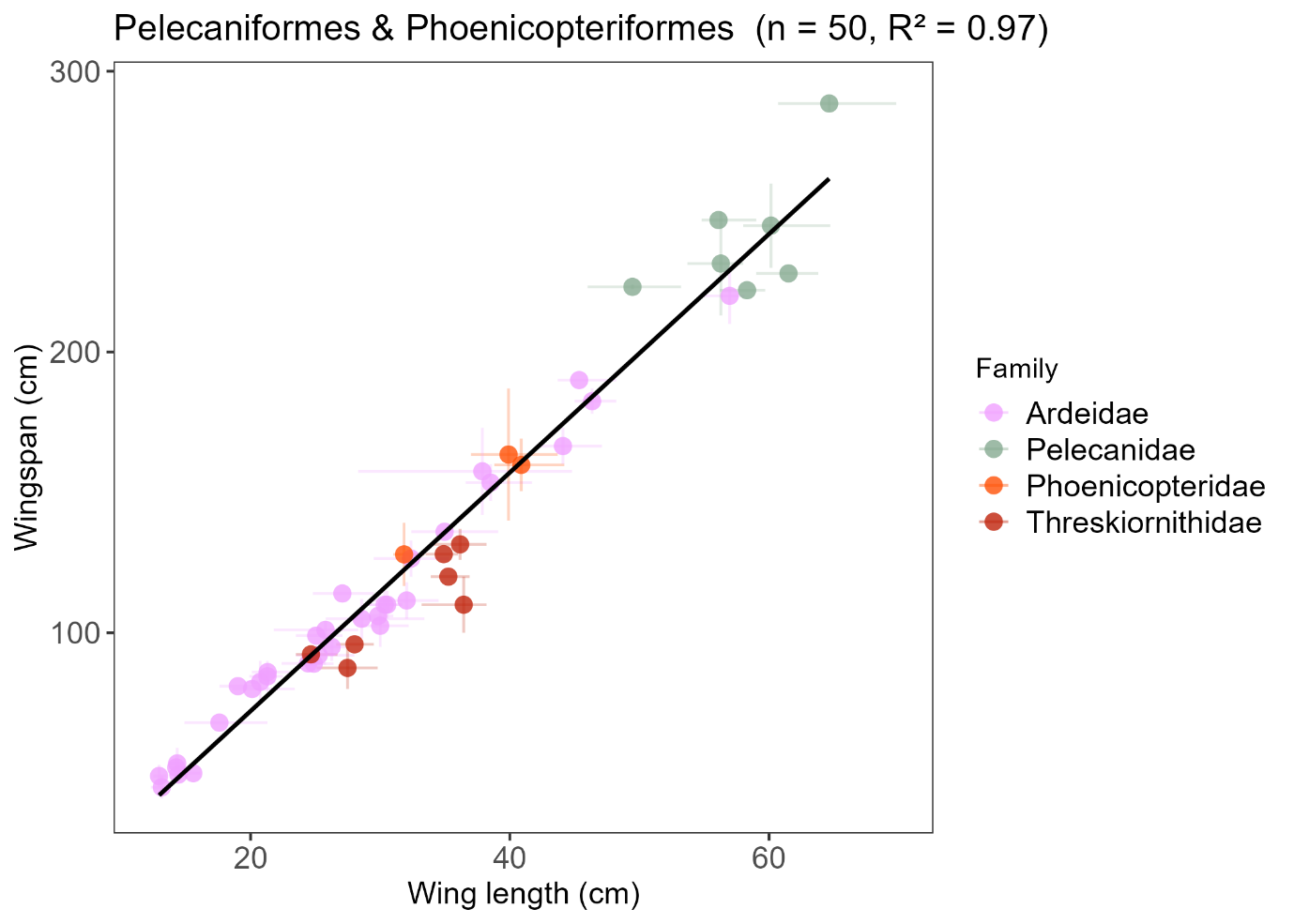

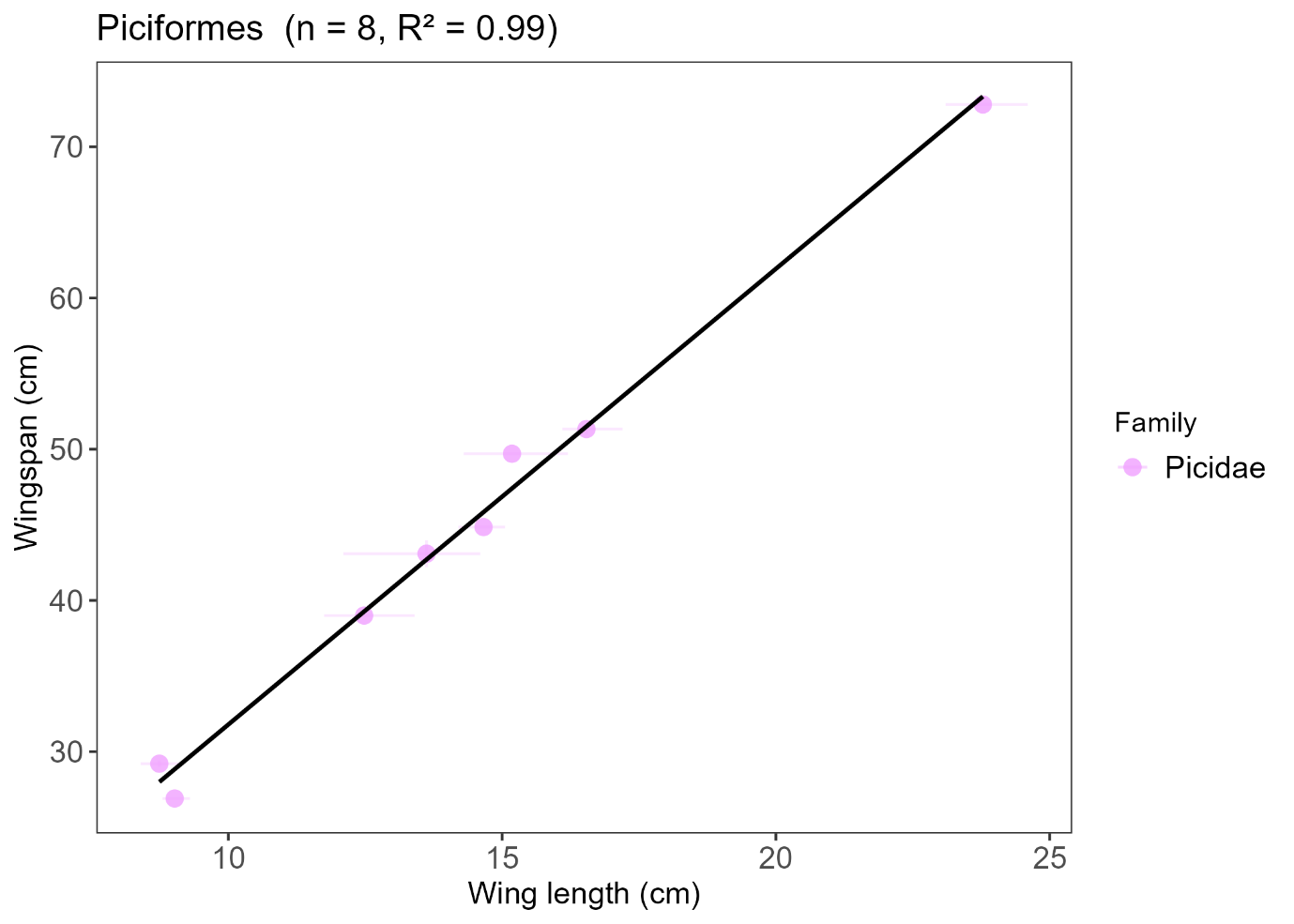

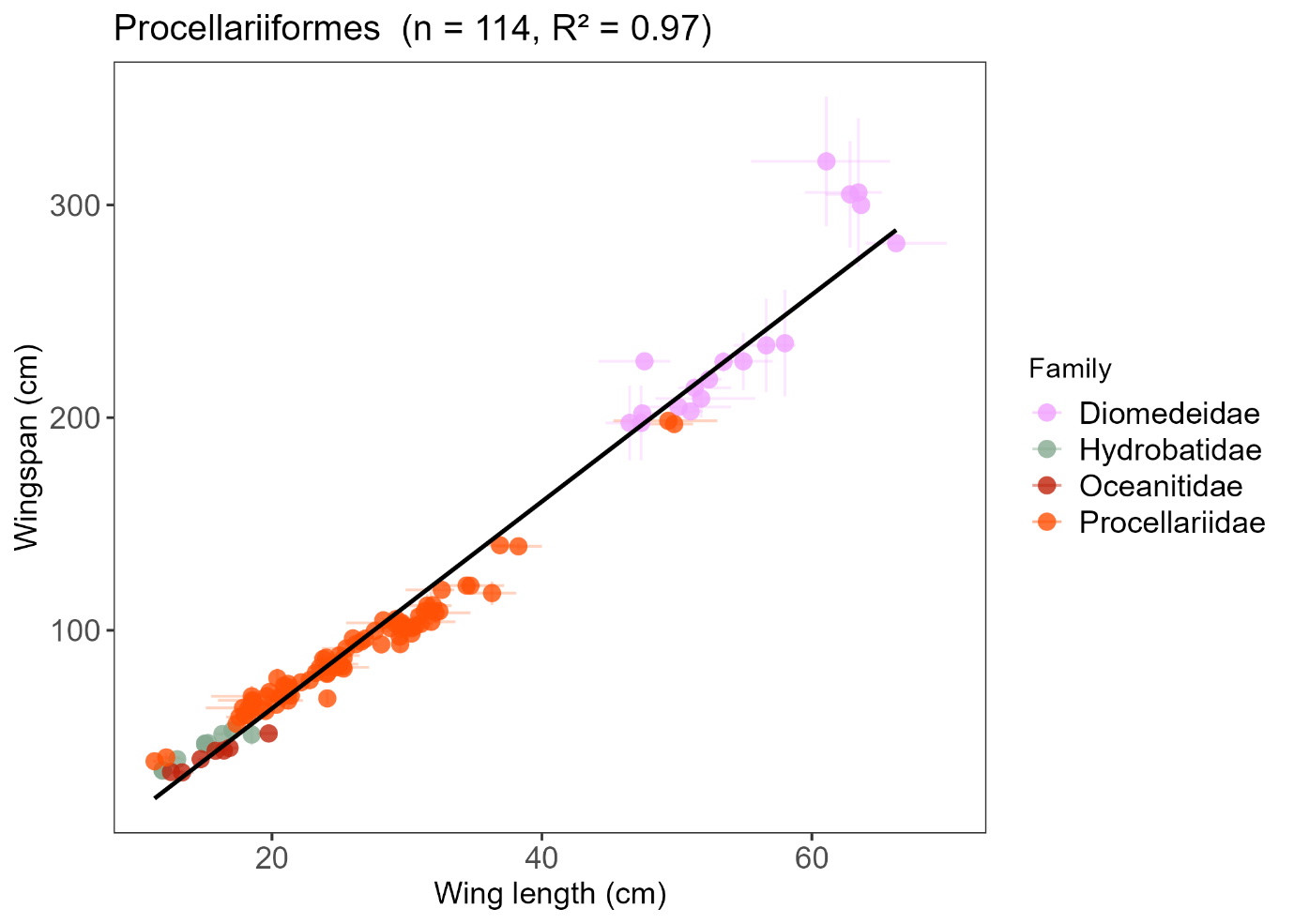

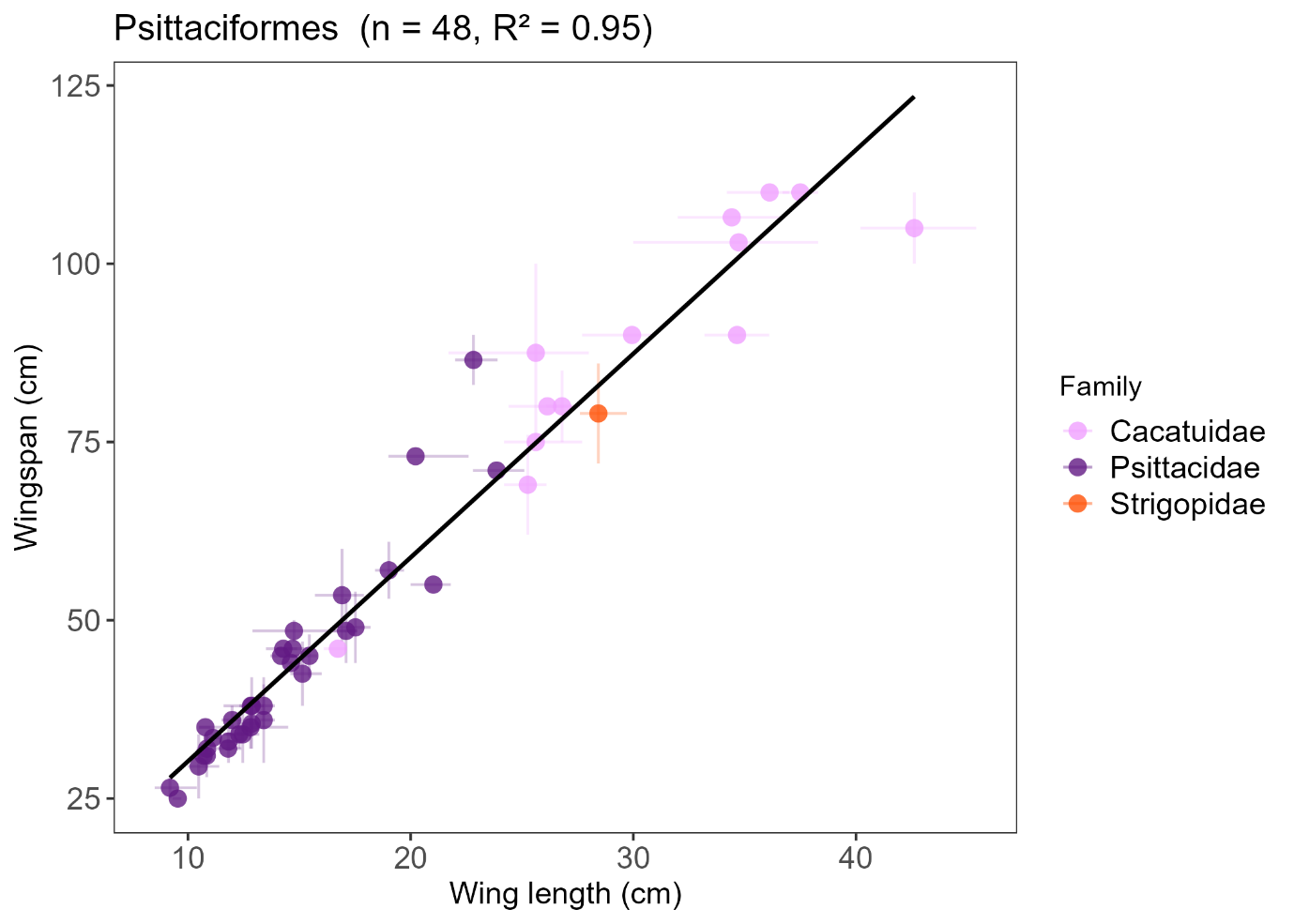

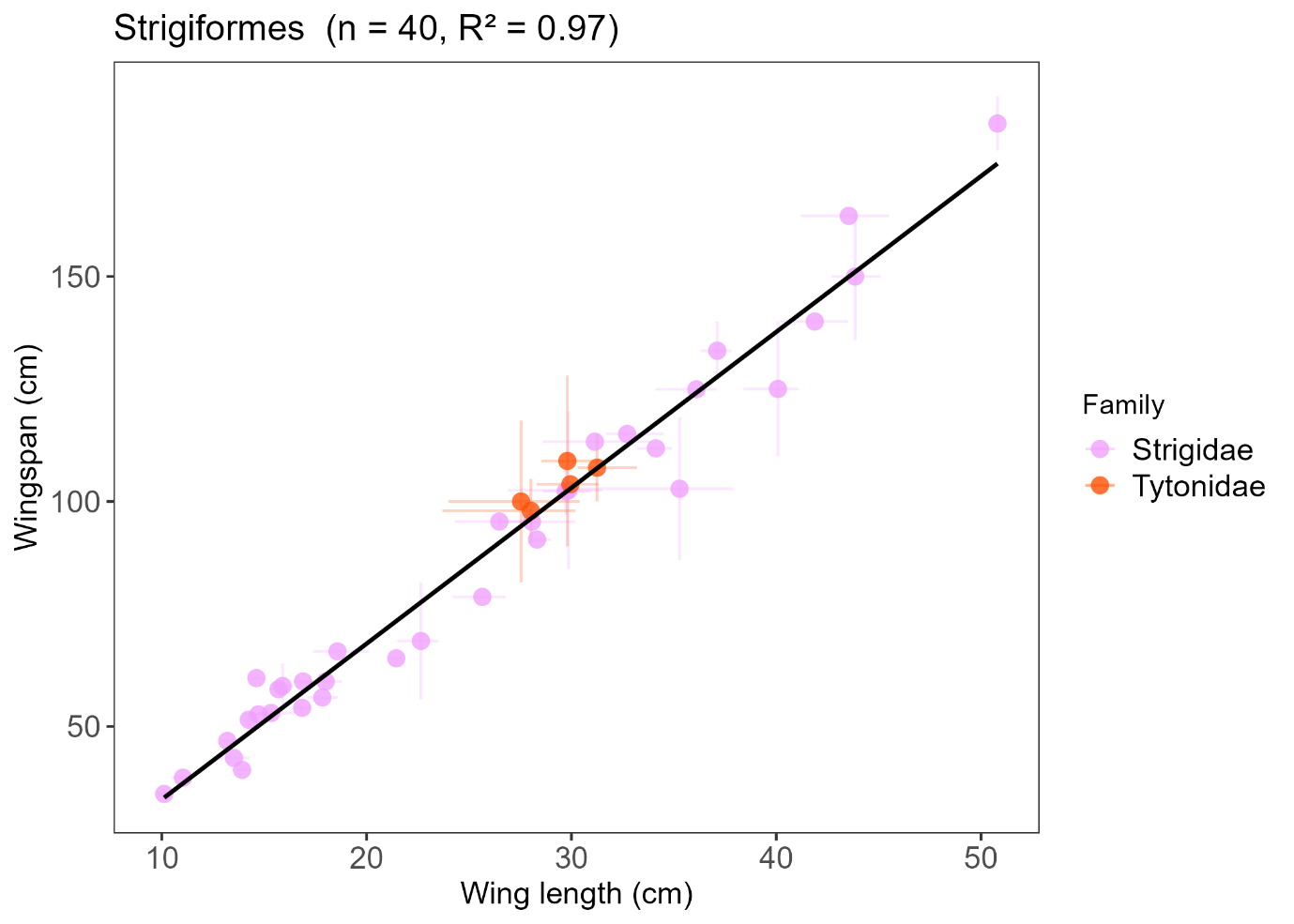

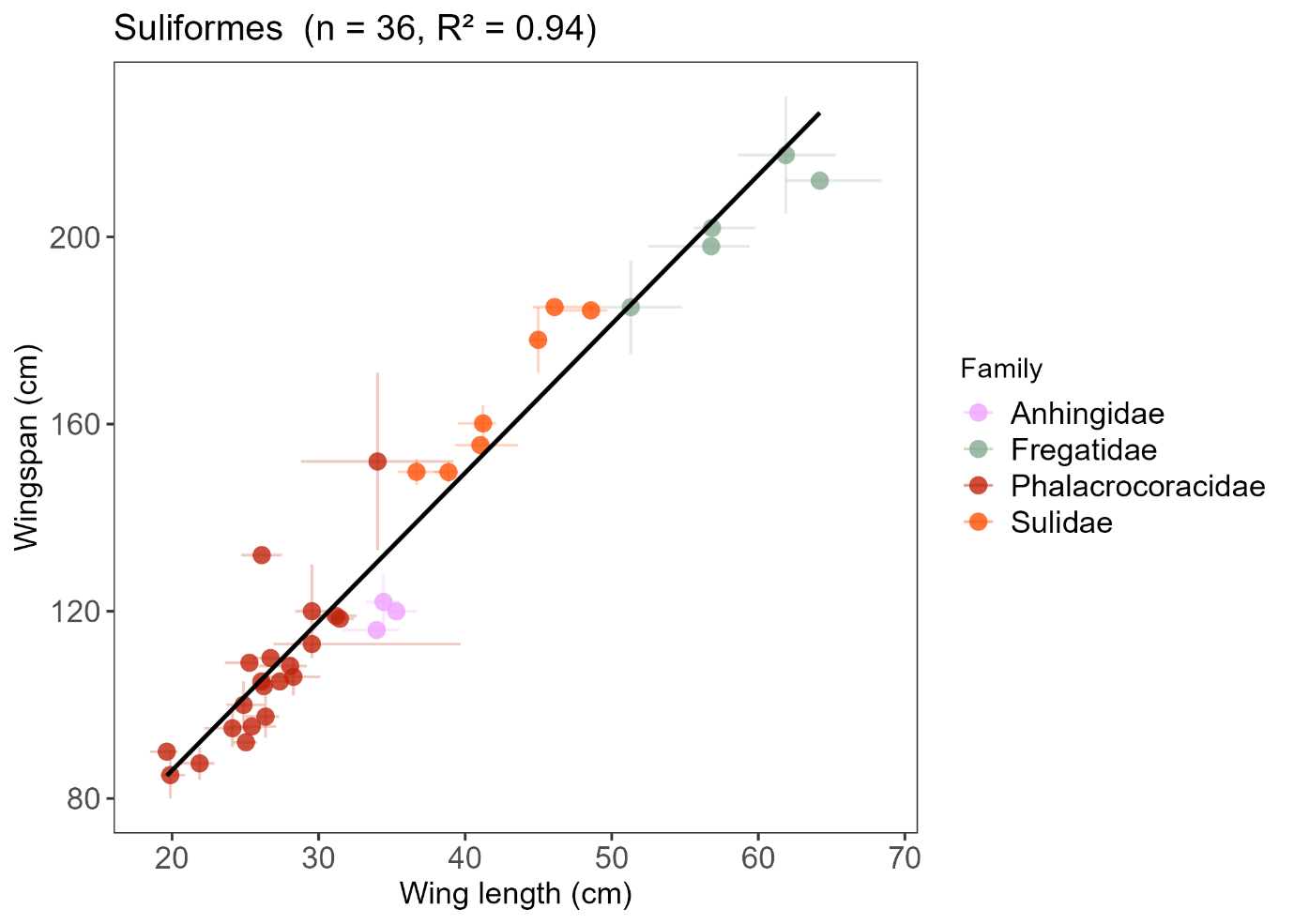
