## Appendix S3 for "Estimating avian wingspan from wing length: an order-level regression approach"

**Appendix S3:** Table S1 and Figure S1

Table S1: Cross-validation (CV) performance of order-level wingspan regression models (1,000 iterations, 20% holdout per order). Percentage prediction error is the absolute difference between the model-predicted and reported midpoint wingspan for each holdout species, expressed as a percentage of the reported midpoint. Median intraspecific range is the intraspecific variation in wingspan for species with empirical range data, expressed as the difference between the reported midpoint and maximum (or minimum) wingspan as a percentage of the midpoint; where median prediction error is less than or equal to the median intraspecific range, the model predicts within the bounds of natural variation. Orders where median prediction error exceeded the median intraspecific range are denoted by *.

| **Order** | **Median prediction error (%)** | **Mean prediction error (%)** | **Predictions within intraspecific range (%)** | **Median intraspecific range (%)** | **Number with intraspecific range** |
| --- | --- | --- | --- | --- | --- |
| Falconiformes | 3.4 | 4 | 87.8 | 7.3 | 46 |
| Psittaciformes | 6.1 | 6.9 | 75.1 | 9.0 | 26 |
| Accipitriformes | 4.6 | 6.0 | 72.7 | 7.8 | 177 |
| Piciformes* | 2.4 | 3.6 | 66.7 | 0.7 | 3 |
| Cuculiformes | 5.1 | 5.7 | 66 | 7.1 | 9 |
| Columbiformes & Pterocliformes | 2.9 | 4.2 | 65.4 | 5.5 | 16 |
| Ciconiiformes* | 6.2 | 7.8 | 61.7 | 6.1 | 7 |
| Passeriformes | 4.6 | 6.4 | 59.3 | 6.4 | 224 |
| Apodiformes | 2.6 | 3.7 | 59.3 | 3.1 | 61 |
| Strigiformes* | 5.4 | 6.2 | 54.5 | 5.1 | 21 |
| Gaviiformes & Podicipediformes* | 4.4 | 4.4 | 50.6 | 3.9 | 4 |
| Pelecaniformes & Phoenicopteriformes* | 6.3 | 7.4 | 48.5 | 5.9 | 29 |
| Anseriformes* | 6.0 | 7.3 | 44.5 | 5.5 | 49 |
| Charadriiformes & Phaethontiformes* | 5.4 | 6.9 | 42.5 | 4.7 | 138 |
| Otidiformes | 8.6 | 11.1 | 38.8 | 11.5 | 5 |
| Suliformes | 4.3 | 5.7 | 35.1 | 4.4 | 16 |
| Coraciiformes* | 7.1 | 8.7 | 32.7 | 3.4 | 15 |
| Gruiformes* | 7.0 | 7.5 | 28 | 5.3 | 21 |
| Procellariiformes* | 7.4 | 9.0 | 23.6 | 4.2 | 65 |
| Caprimulgiformes* | 10.9 | 9.9 | 19 | 5.6 | 8 |

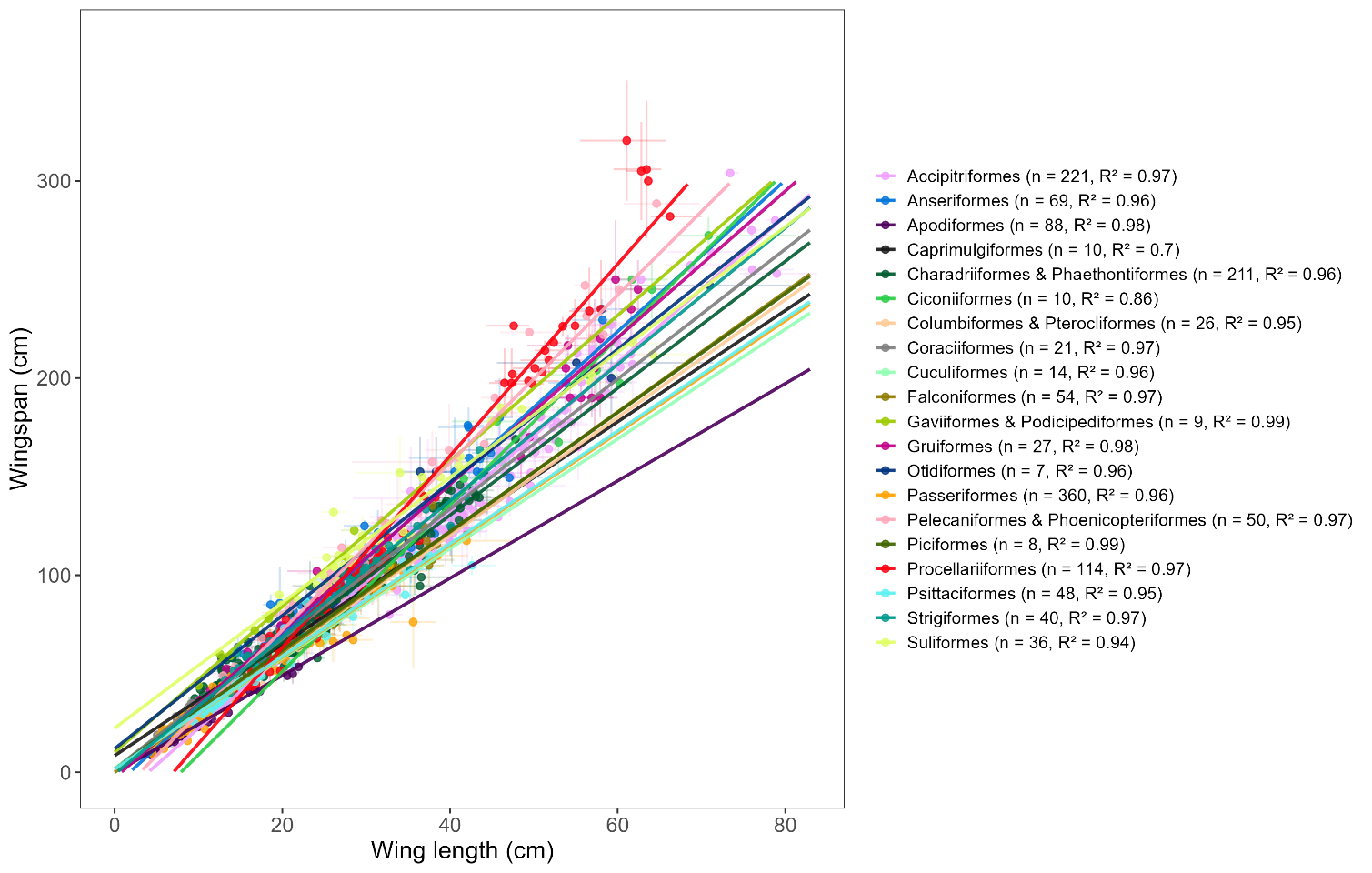

Figure S1: Order-level linear regression models predicting wingspan (cm) from wing length (cm). Each line represents the fitted model for an order-level group, with points coloured corresponding to order-level group. Horizontal and vertical bars represent the range of wing length and wingspan measurements reported for each species, respectively. Model fit (R²) and sample size (n) for each order-level model are provided in the legend.
